# Unraveling Genome- and Immunome-wide Genetic Diversity in Jaguars (*Panthera onca*): Implications for Targeted Conservation

**DOI:** 10.1101/2024.05.06.592690

**Authors:** René Meißner, Sven Winter, Jean Pierre Elbers, Martin Plášil, Ján Futas, Elmira Mohandesan, Muhammad Bilal Sharif, Petr Hořín, Stefan Prost, Pamela A. Burger

## Abstract

Our study examines the declining Jaguar populations in Central and South America, assessing the impact of habitat loss and fragmentation on genetic diversity and local adaptation. We investigated population structure and immunome variability in 25 jaguars to identify unique genetic diversity for informed, targeted conservation. Our genome-wide analyses revealed three distinct geographic populations corresponding to Central America, South American lowland, and South American highland regions. While the highland population displayed lower overall immunome-wide variability, specific innate (Natural killer cell complex, Toll-like receptor) and adaptive (Major histocompatibility complex-class-II) immune genes crucial for adaptive responses showed promising diversity. Nonetheless, South American highland and Central American jaguars are severely threatened. Therefore, we propose re-evaluating evolutionary significant units to prioritize conservation efforts, preserving crucial genetic and adaptive diversity essential for the species’ resilience and long-term survival.

## Introduction

The jaguar (*Panthera onca*) stands out as one of the most remarkable and distinctive members of *Panthera* and is America’s largest feline predator^1^. Unusual for its genus, the jaguar is very muscular for its size and possesses the strongest bite of all extant felids^2^. The species’ remarkable strength is also reflected in its specialized hunting method, which consists of jumping on top of its prey, followed by a fatal bite through the skull^3^. Capable of hunting even the largest mammals, jaguars are not bound to specific habitats, which further consolidates their position as an apex predator^4^. Consequently, they are a keystone species within diverse ecosystems ranging from dense rainforests to open grasslands across Central and South America^5^.

Despite their broad distribution and ecological importance, jaguar populations are highly threatened by deforestation resulting from human population growth and intensifying agriculture^6^. Throughout the Americas, this increasing habitat loss is pressurizing wildlife, but jaguars face additional anthropogenic threats like poaching, persecution and habitat fragmentation, which more severely threatens large carnivores with extensive territories then smaller mammals^7^. For instance, a recent increase in human-made wildfires in Bolivia and Brazil has led to the obliteration of 12% of the Chiquitano Dry Forest^8^, which threatens the resident jaguar population in an already shrinking but important wildlife corridor^9^. Several parts of Brazil already exhibit high levels of habitat fragmentation responsible for local extinction, and reduced gene flow is increasingly threatening the remnant jaguar populations^10^. Furthermore, poaching of jaguars increased drastically in the last years because the traditional medicine market in Asia discovered jaguar fangs as a suitable substitute for decreasingly available tiger fangs^11, 12^. Climate change intensifies these challenges, further increasing habitat degradation, altering prey availability, and escalating the frequency of extreme weather events. However, the jaguar as species is only listed as “Near Threatened” by the IUCN with no current subspecies assignment, although increased regional population decline is acknowledged, particularly in Central America^13^.

A taxonomic re-evaluation of the species based on morphological data^14^ and an extensive microsatellite study^15^ prompted the IUCN to discard the previously assigned subspecies^13^. Because no current subspecies are recognized, the misconception that individual population losses in the jaguar are less severe than in other large carnivores prevails^16^. Yet, populations are increasingly declining, and the identification of evolutionarily significant units (ESUs) in the species is urgent to assist conservation planning and management^17, 18^. When different populations in a species exhibit high degrees of genetic differentiation, ESUs should be assigned to reflect potential adaptational differences in the species subpopulation^19^. Recent studies, using a different set of microsatellites, found noticeable genetic structure within jaguars^20^ and, combined with an initial whole-genome study^21^, supported pronounced distinction between the South and Central American populations. However, to which degree this genetic distinction affects the jaguar’s adaptive potential in its habitats is yet completely unknown.

In general, a species’ adaptability is difficult to be directly assessed because it involves complex interactions between genetic, physiological, ecological, and behavioral factors, which are difficult to measure comprehensively and simultaneously^22^. Nevertheless, several proxies exist that can serve as estimates for an organism’s resilience to environmental changes, and genetic diversity might serve as one of the most reliable estimators because high genetic diversity within a population relates to a higher likelihood of adaptive responses to selective pressures from changing environments^23^. The genetic diversity in immune response genes is directly translated into amino acid sequences, and hence, accessing genetic diversity is more closely tied to the immediate immune function^24^. Increased genetic diversity correlates with the ability to recognize and respond to a broad spectrum of pathogens, therefore, increasing a species’ resilience to diseases^25^.

The recognition of pathogens is essential to activate the innate immune response and specific pattern recognition receptors, such as Toll-like receptors (TLRs) and receptors of the natural killer cell complex, induce this activation^26^. For instance, the Killer cell lectin-like receptors (KLR) of the natural killer cell complex recognize specific molecules on cells, endogenous and non-endogenous, either activating natural killer cells upon ligand binding, or inhibit natural killer cell activity when engaging with self-MHC class I molecules^27^. TLRs, on the other hand, detect a wide array of foreign microbial structures known as pathogen-associated molecular patterns, which can be bacterial cell wall components or virus associated RNA^28^. All these receptors exhibit structural variability and possess high diversity in their ligand recognition domains. They initiate signaling cascades that result in the activation of diverse immune effector mechanisms, including inflammation, and the production of antimicrobial peptides, thereby preventing the establishment and dissemination of pathogens^29^. Averting diseases determines survival, therefore, assessing immunity, especially innate immune response genes, can serve as an indicator of a species’ adaptive potential^30^. Few studies acknowledge this connection and focus on immune response genes and their implications for adaptability, especially in wild felids. While diversity in investigated TLR genes was lower in modern Southern African cheetahs than in African leopards^31^, a comparison of complex immune genomic regions in wild and domestic felids showed a general higher variability in natural killer gene complex (NKC) than in major histocompatibility complex (MHC) encoding genes^32^.

In this study, we combined immune genetic and population genomic approaches to shed light on the sparsely understood population structure of the jaguar using contemporary and historic data. We further compared genome-wide diversity to that of the innate and adaptive immune response genes in light of the observed population structure. Our sampling encompassed large portions of the present-day distribution of the species to assist the re-evaluation of the jaguar’s systematics. We generated whole-genome data for both contemporary and historical individuals, and also incorporated individuals from an already existing dataset^21^. The inclusion of historic samples is essential for understanding the present population structure in an evolutionary context. Furthermore, we examined the immunome, and investigated the diversity of three distinct immune response gene families, TLR, NKC, and MHC class II, in contemporary jaguars. With that, we aimed to enhance our understanding of important immune response genes underlying the species’ disease resilience. Despite habitat degradation impacting genetic diversity and structure in jaguar populations^33^, there has not been any assessment of the species’ immune response genes to evaluate its adaptive potential.

## Results

This study aimed to establishes criteria for the assignment of ESUs in jaguars and to evaluate the species’ adaptive potential by generating comprehensive whole-genome- and immunome-wide data for 11 historical and 14 modern jaguar individuals. Using paired-end sequencing, new short reads for two modern and 11 historical individuals were generated, and pre-existing short-read data from 12 modern individuals from a previous study were utilized^21^. All individuals were assigned to one of the following three jaguar populations based on their geographic origin: samples from Central America were assigned the Central American population, samples from Amazon (-Orinocan) lowland, Gran Chaco, and Pampas belonged to the South American lowland population, and samples from Cerrado, Caatinga, and Atlantic Forest were assigned to the South American highland population. We examined population structure in jaguars using genome- and immunome-wide SNP data, firstly to validate the assumed population structure based on ecoregions, and secondly to investigate the distinction between individual populations. Sufficient genetic distinction is an important criterion to assess ESUs in a species, therefore, we further examined diversity parameters such as nucleotide diversity and heterozygosity and measured pairwise fixation between all jaguar populations. Additionally, to obtain an approximation of the species’ adaptive potential, the genetic diversity of two innate (TLR, NKC) and one adaptive (MHC class II) immune response gene family was examined.

### Genome- and immuno-wide SNP data reveal three distinct jaguar populations

The genome-wide analysis of 689,785 unlinked SNPs from 25 individuals showed that the jaguar can be separated into different geographic populations across its distribution. The PCA revealed a clear separation into three clusters (Figure 1b), which correspond to three distinct regions across Central America, South American lowland and South American highland. The Central American jaguars are separated by PC1 (28.52 %) from the South American individuals, while the two clusters of South American highland and lowland individuals are separated by PC2 (9.05 %) and PC3 (7.26 %). Two South American highland individuals, PO31 and PO32, clustered together with the South American lowland population, while one individual of the South American lowland population, AM404, grouped within the South American highland population. Within the cluster of the South American highland population, two individuals, AF048 and AF052, were distant to all other individuals but grouped close to each other.

**FIGURE 1:**
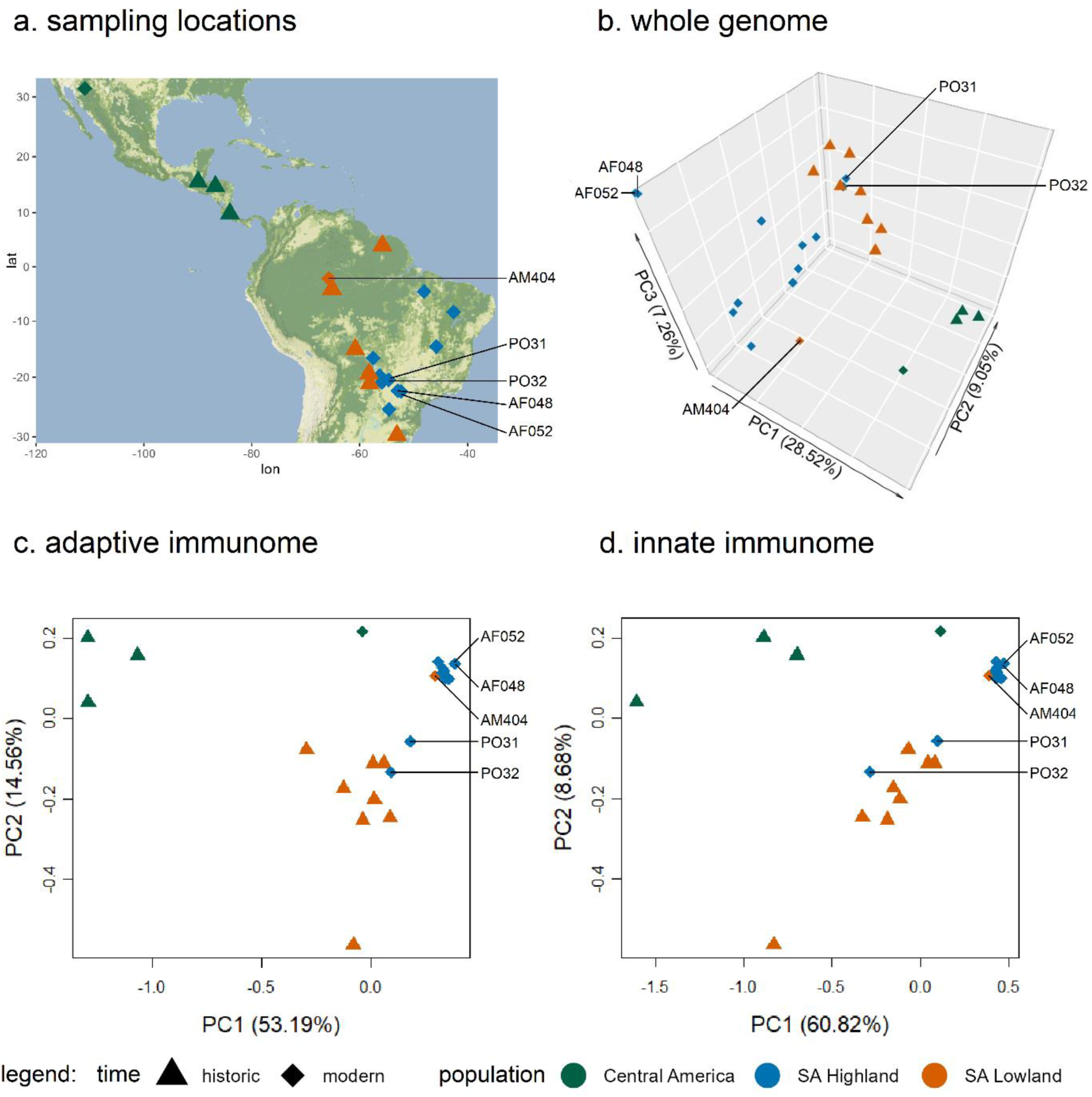
(a) Approximate sampling locations for all 25 jaguar individuals used in this study. (b) Principal component analysis based on 689,785 unlinked genome-wide SNPs showing the first three principal components, (c) principal component analysis of 26,408 unlinked adaptive immunome-wide SNPs, and (d) principal component analysis of 14,155 unlinked innate immunome-wide SNPs showing the first two principal components of 25 jaguar individuals.

Separately conducted PCAs using adaptive (14,155 SNPs), and innate (26,408 SNPs) immunome data generally exhibited less pronounced clustering of jaguar populations compared to the PCA based on genome-wide SNPs, although all three clusters were discernible (Figure 1b-d). Central American jaguar individuals are distinguished from South American jaguars by PC1 (53.19 % adaptive, 60.82 % innate), while PC2 (14.56 % adaptive, 8.68 % innate) separated the South American lowland and highland populations. PC3 of the adaptive and innate immunome explained less than 5 % of the observed variation (Supplementary figure S1). However, unlike the PCA of the whole genome (Figure 1b, Supplementary Figure 2), the within-cluster variation of adaptive and innate immunome was comparatively lower in the South American highland population than in both other populations. Notably, the individuals AF048 and AF052, contrary to their clustering in the PCA based on the whole genome data, did not group distant to other individuals from the South American lowland population. Furthermore, the modern individual MDAZ grouped closer with individuals from the South American highland population in both immunome-wide PCAs than with those individuals from the Central American population and unlike in the PCA based on the whole genome data.

The highest differentiation between populations was observed between Central America and the South Americas highland (F_st_ = 0.179829), while the differentiation between Central America and the South American lowland (F_st_ = 0.072500) and between the South American lowland and highland population (F_st_ = 0.043549) were lower. Considering both South American lowland and highland individuals as belonging to a single population, the differentiation between this unified South American population and the Central America population was slightly lower (F_st_ = 0.133887) than the differentiation between Central America and the South Americas highland.

Using the same unlinked SNPs of the genome-wide PCA, the admixture analysis reflected the previously detected population structure, and revealed limited genetic exchange between the three distinct jaguar populations (Figure 2a). At K = 2, the Central American population separated from both South American populations with some admixture present in the lowland population, while at K = 3, the South American lowland and highland populations segregated. The same individuals (AM404, PO31, PO32) that deviated in the PCAs from their assumed populations based on geographic origin are also evident here. For K > 3, no further population structure is apparent that can be correlated with geographical patterns in jaguars. Noticeably at K = 4, the individuals AF048 and AF052 from the South American highland population, which clustered distant to their assigned population based on origin in the whole genome PCA, grouped with two individuals from the South American lowland population, PO04 and PO07, without any apparent connection to a geographic pattern.

**FIGURE 2:**
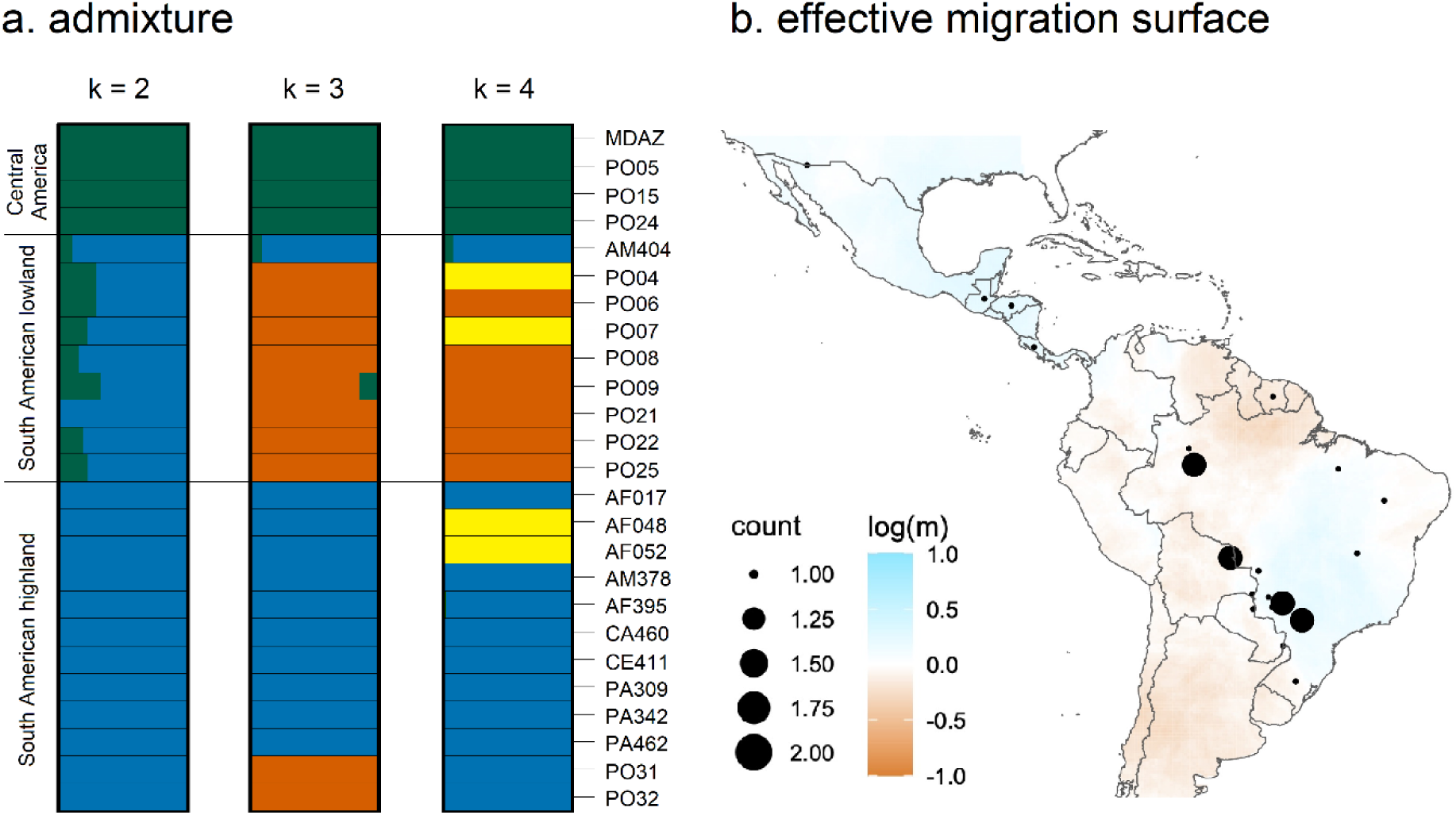
(a) Admixture analysis of 689,785 unlinked genome-wide SNPs with K ranging from 2 to 4. (b) Effective migration surface for all 25 jaguar individuals supporting migration within the Central and South America highland population but not within the South American lowland population. Blue indicates higher than overall average migration, white corresponds to the overall mean migration, and brown indicates strong barriers to migration.

Furthermore, we examined migration between jaguar populations by estimating the effective migration surface for all 25 individuals (Figure 2b). Higher than average migration was observed within the Central American and South American highland population, lower than average migration was observed within the South American lowland population. However, the lower-than-average migration could also be an artifact for the limited sample size with known GPS coordinates for this region.

### Genome-wide and immunome nucleotide diver between jaguar populations

The nucleotide diversity of the three jaguar populations ranged from π = 0.09047 to π = 0.15330 in the immunome and from π = 0.12674 to π = 0.16441 in the whole genome (Figure 3a). For the whole genome, the highest nucleotide diversity (π = 0.15330) was observed in the South American lowland population, while the lowest diversity (π = 0.09047) occurred in the South American highland population. The lowest immunome-wide nucleotide diversity (π = 0.12674) was also observed in the highland population, while the Central American population showed the highest immunome-wide nucleotide diversity (π = 0.16441). For the South American lowland and highland population, genome-wide nucleotide diversity was higher than immunome-wide diversity, and for the Central American population genome-wide nucleotide diversity was lower than immunome-wide diversity. Nucleotide diversity differences between exome, immunome, adaptive immunome, and innate immunome were marginal, except for the Central American population, where the nucleotide diversity of the exome was lower than that of the immunome (Supplementary table 2).

**FIGURE 3:**
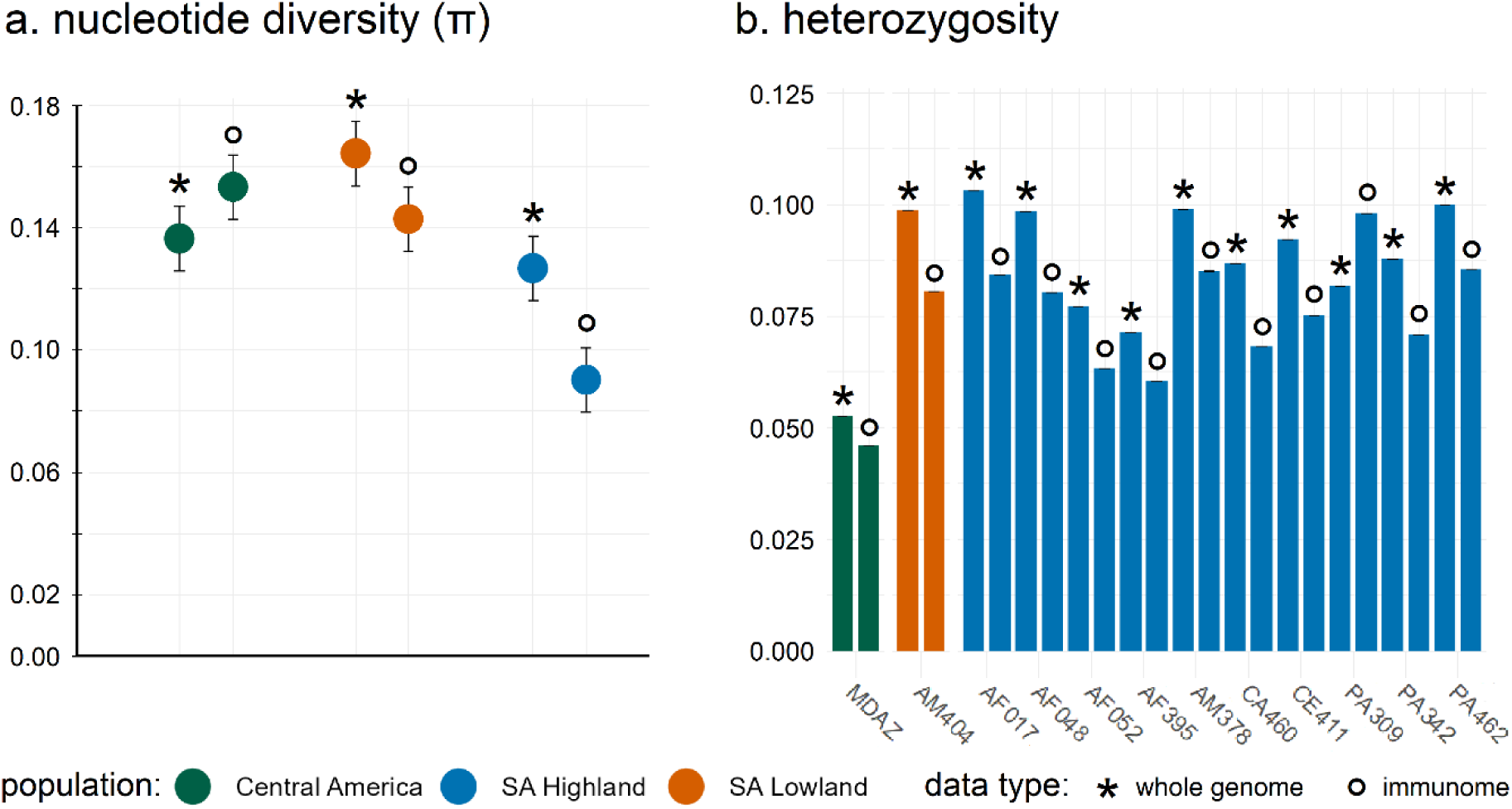
(a) Whole genome- and immunome-wide nucleotide diversity [π] for the three jaguar populations, and (b) individual whole genome- and immunome-wide heterozygosity for modern jaguars (excluding PO31 and PO32). Different population in green = Central America, orange = South American lowland, and blue = South American highland; Data used for analyses either asterisk = whole genome, or circle = immunome.

The heterozygosity of the whole-genome and immunome from 12 modern jaguar individuals (excluding PO31 and PO32) ranged between 0.10 and 0.04 % (Figure 3b). The lowest values were observed in MDAZ from the Central American population (whole genome = 0.053 %, immunome = 0.046 %) and two individuals, AF052 (whole genome = 0.077 %, immunome = 0.063 %) and AF395 (whole genome = 0.071 %, immunome = 0.061 %), from the South American highland population. The genome- (0.053 %) and immunome-wide (0.046 %) heterozygosity of MDAZ was almost halve that of some South American highland individuals, e.g., AF017 (whole genome = 0.103 %) and PA309 (immunome = 0.092 %). Heterozygosity of the immunome was consistently lower than that of the whole genome, except for individual PA309 where the heterozygosity of the immunome was higher than that of the whole genome.

It is worth to note that our sampling did include both modern and historic samples, which could impact genomic inferences. However, we only observed very minor differences when running the analyses only on transversions, as transitions can potentially be impacted by DNA damage in historic samples^34^. This indicates that the results are not driven by damage pattern in the historic samples.

### Specific innate and adaptive immune response gene diversity

We were able to examine nine genes of the NKC, all ten TLR genes expected in felids, and two MHC class II genes in the 12 modern jaguar samples, which showed sufficiently high coverage (> 10x, Table 1) to call heterozygous alleles reliably. These samples included 10 individuals from the South American highland and one sample each of the South American lowland and Central American population, which have been also used for the heterozygosity estimation. The genes belonging to the NKC included KLRA, six KLRC genes: four members of KLRC1 (KLRC1-1, KLRC1-4, KLRC1-5, KLRC1-6), KLRC2, and KLRC3, and KLRH4, and KLRJ. TLR genes included TLR1-10, and MHC class II genes included DMA and DRA. The nuclear allele count ranged between 2-23 and was higher or equal as high compared to the resulting amino acid sequences that ranged from 1-23 (Table 1). Most genes, irrespective of their family, exhibited high heterozygosity (≥ 0.5) and haplotype diversity (≥ 0.6), with only a few genes falling below these values. Notably, TLR3 and TLR8 both exhibited heterozygosity below 0.01 and haplotype diversity below 0.4. KLRA and DMA displayed the lowest number of nuclear alleles (N_alleles_ = 2) and TLR7 and TLR8 showed the lowest number of amino acid sequences (N_amino acid_ = 1). The lowest heterozygosity was observed in TLR3 und TLR8 (H = 0.083), which also showed the lowest haplotype diversity (h = 0.3254). For several genes (TLR2, TLR6, DRA), all jaguar individuals were heterozygous.

**TABLE 1:**
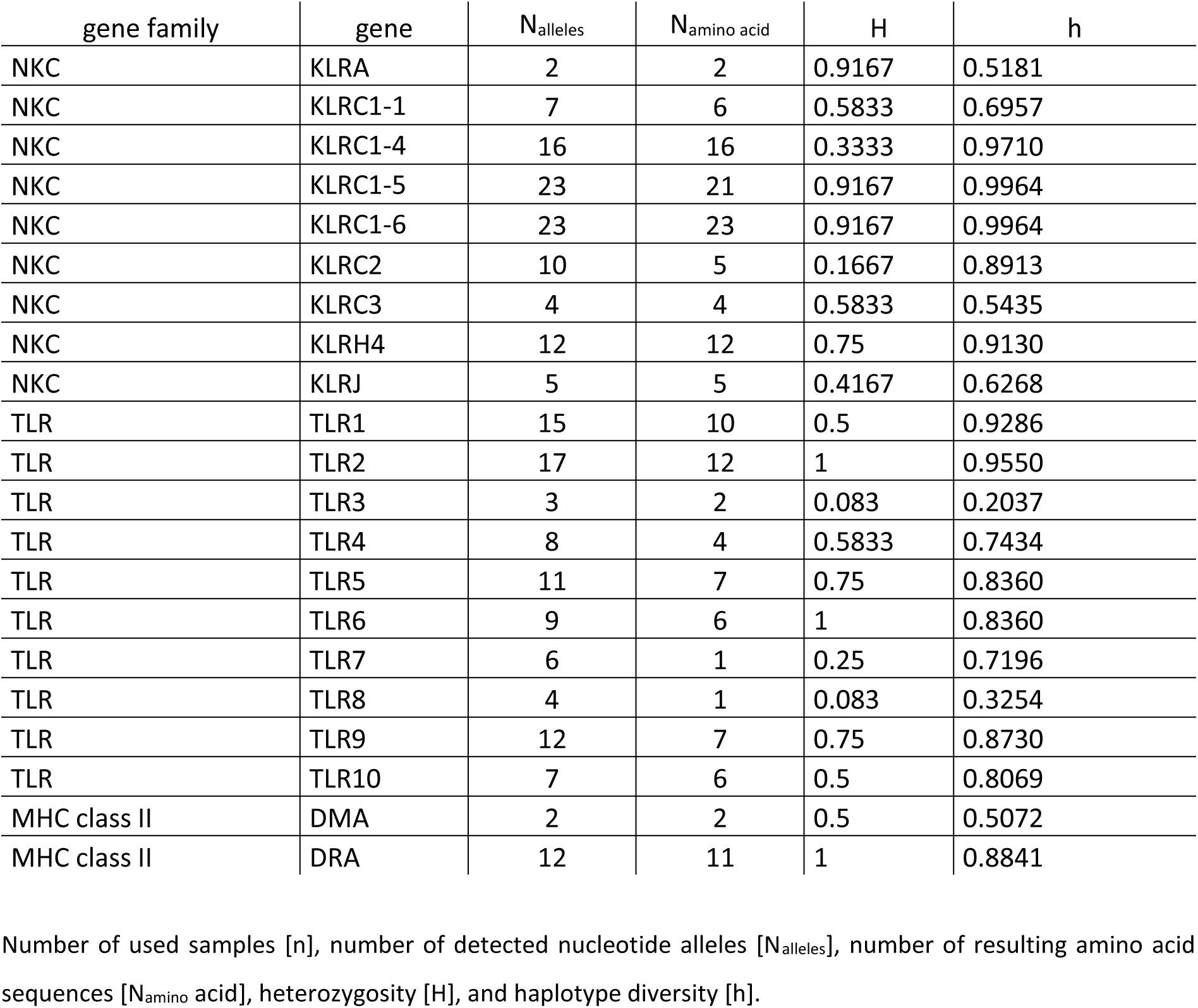
Comparison of innate (NKC, TLR) and adaptive (MHC class II) immune response gene diversity within 12 modern jaguars (excluding PO31 and PO32).

## Discussion

### Population and immune genomic implication for conservation

Species lacking the assignment of ESUs, reflecting their genetic distinction, give the impression of genetic uniformity. A problem potentially leading to oversight of regional population decline that threatens unique genetic diversity^35^. The jaguar is currently without an ESU or subspecies assignment, regardless of its large continent-spanning distribution. Consequently, the whole species is only listed as “Near Threatened” by the IUCN, since regional population decline is not sufficiently reflected in the most recent assignment^13^. However, jaguars are more severely endangered in certain parts of their distribution than whole population size estimations indicate, and its unique genetic diversity might already be threatened in regions of increased population decline.

Earlier microsatellite studies investigating population structure in jaguars did not detect noticeable genetic separation within the species^15^. However, our genome- and immunome-wide analyses support a clear genetic differentiation that corresponds to distinct geographical regions in South and Central America. The separation between Central and South American jaguar populations has already been proposed using modern whole-genome data but lacked the sampling to make a definite statement about delineation in South America^21^. Our genomic data additionally incorporated historical individuals from an expanded sampling area and supported not only the distinction between Central and South America but separated two groups within South America. Furthermore, recent genetic studies employing different sets of microsatellites revealed more population structure in Central American^20, 36^ and South American^37^ jaguars than previously thought.

While microsatellites support a separation into three distinct populations in South America, namely Amazon, Pantanal, and Atlantic Forest, our genomic data only suggested two: a South American lowland population including the rainforest of the Amazon, Pantanal, and Gran Chaco, and a South American highland population including the biomes of the Cerrado, Caatinga, and Atlantic Forest. Contrasting the microsatellite-based separation, whole-genome data did not identify a separate Pantanal population. This could be attributed either to the higher power of genomic SNPs to identify distinct groups in clustering methods, compared to microsatellite data, which tends to overestimate local distinction^38, 39^, or to our limited sampling in the area compared to the microsatellite studies^37^. Overall, the identified genetic structure in jaguars was consistent with earlier studies, independent of the used population genetic markers. Nevertheless, SNPs are more precise than microsatellites in population-level diversity estimation and allow for additional adaptability assessments^38^. Therefore, we only considered three distinct jaguar populations in the following based on whole-genome data, and these populations are Central America, South American lowland, and South American highland.

While most jaguar individuals were correctly assigned to the three populations based on geographic ecoregions, three samples showed conflicting signals between genomic and geographic population assignments. Unlike other individuals from the Amazon, AM404 did not cluster with the South American lowland population but with the South American highland population, which might be explained by limited sampling in the region. So far, no genomic data for individuals from Central South America exist, thus it is still unclear where precisely the boundary lies between the South American lowland and highland populations. Therefore, AM404 could potentially originate from the border of both populations which led to its misclassification. Comparably, the individuals PO31 and PO32 could also be falsely assigned to the South American highland population based on their assumed geographic origin. The exact sampling locality for both individuals is unknown but they originate from the Brazilian state Mato Grosso do Sul. This state encompasses two different ecoregions, Pantanal and Cerrado, which belong either to the South American lowland population or to the South American highland population. As the Cerrado covers most of the state’s area, we initially assigned PO31 and PO32 to the South American highland population based on probability. However, using whole genome data, both individuals cluster with other samples of the South American lowland population, and therefore more likely originate from the Pantanal. To assist sample assignments in the future, extending the sampling area especially in regions where both South American jaguar populations border each other is necessary, and the exact sampling locality must be noted. Yet, geographic ecoregion and genomic data mostly agreed in the assignment of individual samples to the three distinct populations.

Despite their clear separation, jaguars seemed less strongly divergent than other big cats^40, 41^. The highest genetic distinction among jaguars was observed between the Central American and the South American highland populations (F_st_ = 0.179829), which are also the most geographically distant. Conversely, the lowest genetic differentiation was identified between the South American lowland and South American highland populations (F_st_ = 0.043549), which are geographically adjacent to each other. Considering both South American lowland and highland jaguars as part of a single population due to their moderate level of distinction, the differentiation between South and Central America was still high (F_st_ = 0.133887). This further supports the Central American jaguar as the most distinct (similar to Lorenzana et al., 2022), and could justify classifying this population as a separate subspecies^42, 43^. Based on our findings, a re-assessment of any ESU is urgent to acknowledge the species’ genetic distinction and to assist differentiated jaguar conservation efforts^18^.

The Central American and South American highland population represent the northernmost and easternmost jaguar populations and are on the edge of the species’ distribution^16^. Therefore, these populations are more severely threatened by habitat loss because any level of habitat fragmentation is more likely to result in genetic isolation and impoverishment compared to the core population^44^. The largest threat to the jaguar, deforestation for land development purposes, already causes intense habitat fragmentation in Central America^36^ and limited gene flow between populations in the South American highlands^37^.

Furthermore, two highland individuals, AF048 and AF052, already showed increased levels of inbreeding^21^, and at least AF052 also exhibited decreased levels of heterozygosity in our study. Similarly, increased inbreeding was observed for several subpopulations of Central American jaguars^45^, and compared to both South American populations, the Central American population was even more genetically depauperate^20^. Our results further supported these findings, as the single contemporary Central American jaguar in our study, MDAZ, was the least heterozygous individual within all contemporary individuals for which we were able to calculate heterozygosity, including South American lowland and highland populations. Nevertheless, both populations, Central America and South American highland, show reduced genetic diversity, thus their adaptive potential might already be lowered compared to the South American lowland population.

### Immune genetic diversity to infer the adaptive potential in jaguars

To investigate the adaptive potential of the three jaguar populations, we took an in-depth look into the species’ immunome as it represents the entirety of annotated immune genes and is believed to consist of mostly genes under selection^46^. We discovered that the immunome mainly reflected the whole genome’s population structure in jaguars, but the within-population variability was noticeably reduced in the South American highland population. Comparing nucleotide diversity between the adaptive and innate immunome, we did not observe any noticeable differences. However, the nucleotide diversity of the immunome was either lower than that of the whole genome (in South American lowland and highland populations) or higher (in the Central American population). Compared to both South American populations, jaguars of the Central American population occur in regions of high human population density, which results in increased habitat fragmentation and an expanding human wildlife interface^47^. Consequently, Central American jaguars experience a strong population decline^45^, while their interactions with humans, their pets, and livestock increase simultaneously, exposing them to different and new pathogens^48^. Despite the accompanying inbreeding, this high pathogen exposure might favor diversifying selection in immune response genes to maintain disease resilience^49^, while other gene families, which are subjected to lower selection pressure, decrease in diversity^50^. This could explain the higher immunome-wide nucleotide diversity in Central American jaguars compared to the whole genome’s diversity. Therefore, monitoring immune genetic diversity might be a better indicator for adaptability in threatened species than genome-wide diversity^51^. Yet, the overall lowest nucleotide diversity, both in the immunome and whole genome, did not occur in the Central American but rather in the South American highland population.

However, additional investigations of the NKC, TLR, and MHC class II genes in the less genetically variable highland population did not reveal genetic impoverishment. In particular, KLRC1 genes were highly variable and exhibited increased levels of heterozygosity and haplotype diversity compared to other mammals including camels^52^, lemurs^53^, and human^54^. When examining the diversity of NKC and MHC class II genes within the Central and South American jaguar individuals, we observed that NKC variability surpassed that of MHC class II genes. This aligns with prior research conducted in domestic cats, where NKC exhibited greater variability compared to MHC, encompassing class I, II, and III genes^32^. The number of MHC class II DRA alleles (n = 12) found in the 12 modern jaguars in this study was proportionally higher than the allelic diversity (n = 13) in DRB genes from 46 modern and historic African and Asiatic cheetahs^40^. Yet, previous studies on Namibian cheetahs did not find evidence of a compromised immunocompetence even with low allelic diversity (n = 4 in 94 individuals) in MHC class II DRB genes^55^. Therefore, MHC gene diversity might be a poor indicator of population health in some wild felids. It is interesting to note that also ungulates such as camels exhibit a similarly low diversity in DRA (n = 3) and DRB (n = 5) alleles^56^ without known reduction in their immunocompetence^57^. Furthermore, the diversity of TLR genes in the jaguar was high compared to those in the cheetah and African leopard^31^, suggesting that jaguars might possess a heightened innate immunity, which could compensate for potential genetic depletion in certain adaptive immune genes^58^.

Due to insufficient genome coverage, we were unable to adequately investigate the selected gene families within the Central American and South American lowland populations. Consequently, we could not contextualize the observed genetic diversity in the South American highland population within a broader framework. Missing a within-species comparison, the information gathered from the three gene families is limited and the immunome-wide variability might be a better approximation for the jaguar’s adaptability until future studies provide sophisticated distribution-wide data on these specific genes. Nonetheless, the immunome of the South American highland population was less variable than that of both other populations, which might already indicate a reduced adaptive potential, and stressed the urgency to further protect this jaguar population.

## Conclusion

Our study highlighted geographic differentiation within jaguar populations to address questions concerning distinct genetic diversity and future adaptive capacities. In our investigation, we have identified noticeable separation within the species and provided insight into potential geographical differences in the immunome among the three identified populations. This underscores the importance of acknowledging and conserving geographic regions showing threatened genetic diversity and habitat-specific adaptations crucial for the jaguar’s future adaptability. Given the accepted significance of ESUs in conservation, we propose a re-assignment of the jaguar to facilitate targeted conservation efforts and propose the Central American population as a potential subspecies. Additionally, it is imperative to identify and protect distinct jaguar populations harboring unique genetic diversity to prevent the loss of potential ecotypes, which could diminish the resilience and evolutionary potential of the entire species^59^. Conservation efforts must therefore take genetic diversity, especially that of immune response genes, and population structure into account to minimize genetic erosion and maximize adaptive potential^60^. Integrating population genomics and immune genetics into conservation strategies is essential for preserving genetic diversity, securing adaptability, and ensuring the long-term survival of jaguars and their corresponding ecosystems, ultimately safeguarding ecosystem function and human well-being^61^.

## Methods

### Sampling

In this study, we used a combined set of 11 historic and 14 modern samples to investigate the population structure of jaguar. Modern samples, collected after 1994, spanned a period of 30 years preceding this study, while historic samples dated back from 1853 to 1990. The data set generated in this study included two modern samples from the Brazilian state Mato Grosso do Sul and 11 historic samples from museum collections in Germany (Natural History Museums of Berlin, State Museum of Natural History Stuttgart, Senckenberg Naturmuseum, Alexander Koenig Research Museum), including three samples from Bolivia, two from the Brazilian Amazonas and one each from Mato Grosso do Sul, Costa Rica, Guatemala, Honduras, Paraguay, and Surinam (Supplementary table 1, Figure 1a). Additional, short-read data for 11 modern samples from Brazil and 1 sample from Mexico were obtained from a previous study^21^ (NCBI BioProject: PRJNA348348). However, the data for an additional Guatemalan sample (MFGT) from the same study was not publicly available at the time of this study.

Based on the level I ecoregions of Central and South America^62^, we categorized the samples to three populations based on the sample’s geographic origin: samples originating from North and Central America were assigned to the Central American population. Samples from the Amazonian (- Orinocan) lowland, Gran Chao, and Pampas were assigned to the South American lowland population, and samples from the Eastern highlands, including Cerrado, Caatinga, and Atlantic Forest, were considered members of the South American highland population. For samples PO31 and PO32, precise geographical locations were not available; only the Brazilian state of Mato Grosso do Sul was specified, and as this state predominantly comprises the Cerrado ecoregion, both individuals were assigned to the South American highland population.

### DNA extraction

Genomic DNA was extracted from jaguar samples including dried skins and bones from museum collections, using a modified salting-out DNA extraction method^63^. Overnight, all samples were rehydrated in nuclease-free water to remove and dilute potential secondary preservatives before DNA extraction. Samples with extremely poor DNA preservation (two bones and three dried skins with total DNA quality per sample < 20 ng) were processed again in the ancient DNA laboratory at the University of Vienna (Austria), following the standard contamination precautions^64^. For bones, ∼200 mg of samples was radiated for 30 min under UV-light and powdered using Retsch MM 400 ball mill (Retsch GmbH, Haan, North Rhine Westphalia, Germany). ∼50 mg powdered sample was used for DNA extraction, using the protocol described in Dabney eta al. (2013). For dried skins, ∼ 1cm x 1cm pieces were first UV-radiated for 30 min and cleaned with 70 % ethanol solution to remove any preservative material. The samples were soaked in 10 ml bleach solution (0.5 % sodium hypochlorite) for 2 min and after washing with ddH_2_O, DNA was extracted using the protocol described in Gilbert et al. (2007)^65^, with slight modifications. The digestion process was done for 48 hours and additional 400 µl of proteinase K (15 mg/ml) was added after ∼24 hours of digestion. All extraction batches included blanks to monitor cross-contamination during sample preparation, and extraction.

### Library preparation & sequencing

Illumina sequencing libraries for good quality samples (total DNA amount > 20 ng) were prepared with the standard NEBNext^®^Ultra™ II DNA Library Prep Kit (New England Biolabs, Ipswich, Massachusetts, USA), while double stranded libraries for the samples extracted in the ancient DNA laboratory were prepared following the Meyer and Kircher protocol^67^. All libraries underwent paired-end sequencing on a Novaseq6000 platform (Illumina, San Diego, California, USA) at Novogene (Novogene company limited, Cambridge, UK).

### Genomic data processing

Detailed command for the analyses can be found in the supplementary file. The short-read data (50-150 bp) of all individuals were trimmed using fastp v.0.20.1 (RRID: SCR_016962^68^) with base correction and low complexity filter enabled to remove sequencing adaptors and polyG stretches at the end of the reads. We used a four bp sliding window to detect regions of poor quality (Phred score <15). Reads were removed if they fit into one of the following categories: reads below 36 bp length, reads with >40% low-quality bases, or reads with five or more undetermined bases (Ns).

The trimmed reads of each of the samples were mapped to the chromosome-level jaguar assembly (GCA_028533385.1^69^) using bwa-mem v.0.7.17 (RRID: SCR_010910^70^). The resulting mapping files were then sorted by assembly position, converted to BAM format, and indexed using SAMtools v.1.9 (RRID: SCR_002105^71^). Duplicate reads were identified with MarkDuplicates in Picard v.3.1.1 (RRID:SCR_006525, Broad Institute, Cambridge, Massetshusetts, USA), followed by INDEL realignment using the GATK v.3.8.1 (RRID:SCR_001876^72^) tools RealignerTargetCreator to identify target intervals and IndelRealigner to perform local realignment. After realignment, we removed all reads from the BAM files that were marked as either unmapped, secondary, QC failed, duplicate, or supplementary, using SAMtools, keeping only reads in proper pairs mapped to non-repetitive autosomal regions.

To identify the repetitive regions in the reference genome, we first masked all known repeats for ‘Felidae’ using RepeatMasker v.4.1.4 (RRID:SCR_012954^73^). Next, we generated a *de novo* repeat library with RepeatModeler v.2.13 (RRID:SCR_015027^73^), followed by another round of RepeatMasker to mask the remaining repeats. The masked regions of both RepeatMasker runs were combined to generate a regions file (BED) of non-repetitive regions on the 18 autosomes in BEDtools v.2.31.0 (RRID:SCR_006646^74^), which was used in the filtering of the BAM files.

### *Inferring population structure &* estimating effective migration surface (*EEMS)*

SNPs were called using ANGSD v.0.940^75^ (flags: -GL 1 -doGlf 2 -doMajorMinor 1 -doMaf 2 -SNP_pval 1e-6 -minQ 20 -noTrans 0 -only_proper_pairs 0 -minInd [75 %] -setMinDepth [5x] -doCounts 1) and further pruned for linkage disequilibrium (LD) with ngsLD v1.1.1^76^. LD was estimated as r2 values for all SNP pairs up to 500 kbp apart. An LD decay curve was plotted for a random sample of 0.05% of all estimated r2 values, with a bin size of 250, to establish suitable thresholds for linkage pruning. All sites were pruned assuming a maximum distance of 75 kbp between SNPs and r2 ≥ 0.1.

A covariance matrix was calculated from genotype likelihoods of 689,785 LD pruned SNPs using PCAngsd v.0.9757^77^ and used to perform a principal component analysis (PCA) with the default settings of the ‘prcomp’ function^78^ in R v.3.6.0^79^. Additionally, the pairwise fixation index (F_st_) for each jaguar population was calculated based on the LD pruned SNPs using the realFSF function of ANGSD. The signatures of admixture between the different jaguar populations were calculated using ngsAdmix v.31 (RRID:SCR_003208^80^). We performed 100 replicates for all ngsAdmix runs ranging from k = 2 to k = 6. We summarized and visualized the results using CLUMPAK^81^. We assessed the model fit of each K value to the data using evalAdmix v.0.95^82^.

Additionally, we inferred effective migration rates between the different jaguar populations performing an EEMS analysis^83^. We performed three replicates with 1 million MCMC iterations each, with a burn in of 100,000, and 7,000 demes. For some of the 25 Jaguar samples used in this analysis, precise coordinates of the collection sites were not available. Consequently, we either assigned coordinates within the wider known sampling area or, in instances where sampling localities were unknown, utilized the coordinates of the center of the respective country or state.

### Estimation of heterozygosity of modern jaguar samples

Genome-wide heterozygosity was estimated for all samples with adequate coverage (>6x) based on the folded side frequency spectrum (SFS). The per-sample site allele frequencies were estimated with ANGSD v.0.940 (flag: -doSaf 1) using the reference genome as ancestral. BAQ computation and mapping quality adjustment were enabled. A minimum score of 30 was set for both mapping and base qualities, and a maximum depth cut-off was set to the 95th percentile of the sample’s depth distribution. The per sample folded SFSs were generated in realSFS (flag -fold 1) with 100 bootstrap replicates. Heterozygosity was then calculated in R v.3.6.0. as the percentage of heterozygous sites out of the total number of sites.

### Nucleotide diversity of 11 historic as well 14 modern jaguars

To calculate nucleotide diversity, a vcf file was generated from genotype likelihoods of the LD-pruned SNPs with ANGSD v.0.930 (flags: -GL 1 -doGlf 2 -doMajorMinor 1 -dovcf 1 -doPost 1 -doMaf 1 --ignore-RG 0 -doGeno 1 -SNP_pval 1e-6 -minQ 20 -noTrans 0 -only_proper_pairs 0 -minInd [75 %] - setMinDepth [5x] -doCounts 1), and diversity statistics were calculated with the stacks populations tool v.1.32^84^ for each population (Central American vs. South American lowland vs. South American highland) separately.

### Immune response gene analyses

To explore the immune genetic diversity and understand the adaptive potential of the central, high- and lowland jaguar populations, we looked at single-copy genes from one adaptive (MHC class II) and two innate (TLR, NKC) immune response gene families. We mapped the short reads to the reference sequences of these genes derived from the jaguar assembly (GCF_028533385.1) using bwa-mem v.0.7.17 and removed PCR and optical duplicates with Picard v.3.1.1. Unmapped reads were removed, and the reads in the resulting mapping file were converted to a new fastq file using SAMtools view v.1.9. The filtered reads were then mapped a second time using bowtie2 v2.4.5 (RRID:SCR_016368^85^) without any clipping (flags: -end-to-end -x -S) to avoid miss-called SNPs due to over-representation caused by falsely clipped reads. The variant calling of both alleles was performed with freebayes v.1.3.7 (RRID:SCR_010761^86^, flags: --report-monomorphic --skip-coverage 10), and written into a vcf file. We used bcftools consensus (RRID:SCR_005227^87^, flags: -H I) to create a consensus fasta file containing variable sites as ambiguity codes. The script used for mapping and variant calling is provided in the Supplementary material. Insertions and deletions (indels) were curated manually, and each called SNP was re-checked for validity using Tablet v1.21.02.08^88^. A SNP was considered valid if at least six independent reads covered the position of concern, and the minor allele was called if at least 33% of all reads supported the position.

We used the PHASE function implemented in DnaSP v.6.12.03^89^ to derive the alleles of the short-read consensus sequences with a threshold of 0.6, allowing for recombination. Heterozygosity [H], and haplotype diversity [h] were calculated in DnaSP using phased allele sequences.

Furthermore, we used rGO2TR^90^ to extract exon regions from the annotation of the jaguar reference genome according to the GO-terms associated with immune response (GO:0006955), adaptive immune response (GO:0002250), and innate immune response (GO:0045087), respectively. To extract the exons, we first conducted a blastp v.2.15.0 (RRID:SCR_004870^91^) search against the Swiss/UniProt database v.2024.02^92^. We then used the BED-files generated by rGO2TR to filter the whole genome BAM files in BEDtool v.2.31.0^93^ to keep only reads mapping in the respective target regions.

## Author contributions

RM, SP and PB designed the project. RM, SW, SP and MBS performed research. PB supervised the immunological and SP the conservation analyses. JF annotated NKC genes in genome assemblies of jaguar and cheetah. RM, SW, SP, JPE, and MP analyzed data. The first draft of the manuscript was written by RM and SW, which was edited by SP, PB, PH and EM. All authors revised the manuscript and approved its final version.

## Acknowledgements

We thank the curators of the Natural History Museums of Berlin, the State Museum of Natural History Stuttgart, the Senckenberg Naturmuseum, and the Alexander Koenig Research Museum. RM and SW acknowledges funding from the Central European Science Partnership (CEUS) project Austrian Science Fund (FWF) I5081-B/ GACR Czech Republic 21-28637 L awarded to PH and PAB. SP is funded by the University of Oulu and the Academy of Finland Profi6 336449 programme “Biodiverse Anthropocenes”. A subset of molecular data generation has been performed in the state-of-the-art ancient DNA facility at the Department of Evolutionary Anthropology, University of Vienna, Austria.

## Data Availability

All data generated and/or analyzed during this study are included within this article’s Supplementary material as well as the detailed list of commands used to generate the presented data and related analyses. The raw sequencing reads are deposited as FASTQ files to NCBI (PRJNA1105330). The immune response gene alignments for 13 modern jaguars are publicly available on the Phaidra repository (https://phaidra.vetmeduni.ac.at/o:2917, https://doi.org/10.34876/cmsy-dd57).

## Supplementary material

**SUPPLEMENTARY TABLE 1:** Individual samples used in this study including sample ID, assigned time period, collection data, country of origin, collection locality, assigned coordinates, and Institution of origin.

| ID | Time | Date | Country | Region | Lat. | Long. | Institution |
| --- | --- | --- | --- | --- | --- | --- | --- |
| AF017 | modern | 1998 | Brazil | Mato Grosso do Sul | -22.358 | -52.928 | Lorenzana et al., 2022 |
| AF048 | modern | 2004 | Brazil | Paraña | -22.492 | -52.336 | Lorenzana et al., 2022 |
| AF052 | modern | 2000 | Brazil | Paraña | -22.492 | -52.336 | Lorenzana et al., 2022 |
| AF395 | modern | 2010 | Paraguay | / | -25.518 | -54.571 | Lorenzana et al., 2022 |
| AM378 | modern | 2008 | Brazil | Pará | -4.574 | -48.053 | Lorenzana et al., 2022 |
| AM404 | modern | 2010 | Brazil | Amazonas | -2.166 | -65.732 | Lorenzana et al., 2022 |
| CA460 | modern | 2004 | Brazil | Piauí | -8.333 | -42.593 | Lorenzana et al., 2022 |
| CE411 | modern | 2011 | Brazil | Bahia | -14.582 | -45.804 | Lorenzana et al., 2022 |
| MDAZ | modern | 2009 | Guatemala | / | 31.354 | -110.934 | Lorenzana et al., 2022 |
| PA309 | modern | 2004 | Brazil | Mato Grosso do Sul | -20.936 | -55.890 | Lorenzana et al., 2022 |
| PA342 | modern | 2003 | Brazil | Mato Grosso do Sul | -19.754 | -56.315 | Lorenzana et al., 2022 |
| PA462 | modern | 2014 | Brazil | Mato Grosso do Sul | -16.649 | -57.499 | Lorenzana et al., 2022 |
| PO04 | historic | 1926 | Brazil | Amazonas | -4.105 | -65.124 | Senckenberg Naturmuseum |
| PO05 | historic | 1953 | Honduras | / | 14.739 | -86.739 | Senckenberg Naturmuseum |
| PO06 | historic | 1988 | Bolivia | Northeast | -15.094 | -60.814 | Senckenberg Naturmuseum |
| PO07 | historic | 1989 | Brazil | Amazon? | -4.105 | -65.124 | Senckenberg Naturmuseum |
| PO08 | historic | 1990 | Paraguay | North | -21.150 | -58.125 | Senckenberg Naturmuseum |
| PO09 | historic | 1853 | Suriname | / | 3.971 | -55.783 | State Museum of Natural History Stuttgart |
| PO15 | historic | before 1900 | Costa Rica | / | 9.819 | -84.053 | Natural History Museum Berlin |
| PO21 | historic | ca. 1900 | Bolivia | East | -19.398 | -58.256 | Natural History Museum Berlin |
| PO22 | historic | 1908 | Bolivia | Rio Grande do Sul | -15.094 | -60.814 | Natural History Museum Berlin |
| PO24 | historic | before 1920 | Guatemala | / | 15.560 | -89.945 | Natural History Museum Berlin |
| PO25 | historic | before 1920 | Brazil | Rio Grande do Sul | -29.746 | -53.091 | Natural History Museum Berlin |
| PO31 | modern | 2001 | Brazil | Mato Grosso do Sul | -20.474 | -54.643 | Alexander Koenig Zoological Research Museum |
| PO32 | modern | 2001 | Brazil | Mato Grosso do Sul | -20.474 | -54.643 | Alexander Koenig Zoological Research Museum |

**SUPPLEMENTARY TABLE 2:** Nucleotide diversity [π] for the whole genome, exome, immunome, adaptive immunome, and innate immunome of all three jaguar populations including standard deviation [σ2] and standard error [σM].

| CENTRAL AMERICA | $\pi$ | $\sigma_2$ | $\sigma_M$ |
| --- | --- | --- | --- |
| whole genome | 0.13673 | 0.03842 | 0.00009 |
| exome | 0.10812 | 0.02899 | 0.00016 |
| immunome complete | 0.15330 | 0.02880 | 0.00072 |
| immunome adaptive | 0.14968 | 0.02872 | 0.00139 |
| immunome innate | 0.15440 | 0.02857 | 0.00102 |
| SA LOWLAND | $\pi$ | $\sigma_2$ | $\sigma_M$ |
| whole genome | 0.16441 | 0.01903 | 0.00006 |
| exome | 0.14557 | 0.01334 | 0.00011 |
| immunome complete | 0.14293 | 0.01463 | 0.00052 |
| immunome adaptive | 0.14075 | 0.01443 | 0.00099 |
| immunome innate | 0.14296 | 0.01473 | 0.00073 |
| SA HIGHLAND | $\pi$ | $\sigma_2$ | $\sigma_M$ |
| whole genome | 0.12674 | 0.02506 | 0.00007 |
| exome | 0.10044 | 0.02242 | 0.00014 |
| immunome complete | 0.09047 | 0.01908 | 0.00059 |
| immunome adaptive | 0.09121 | 0.01912 | 0.00114 |
| immunome innate | 0.09072 | 0.01911 | 0.00083 |

**SUPPLEMENTARY FIGURE S1:**
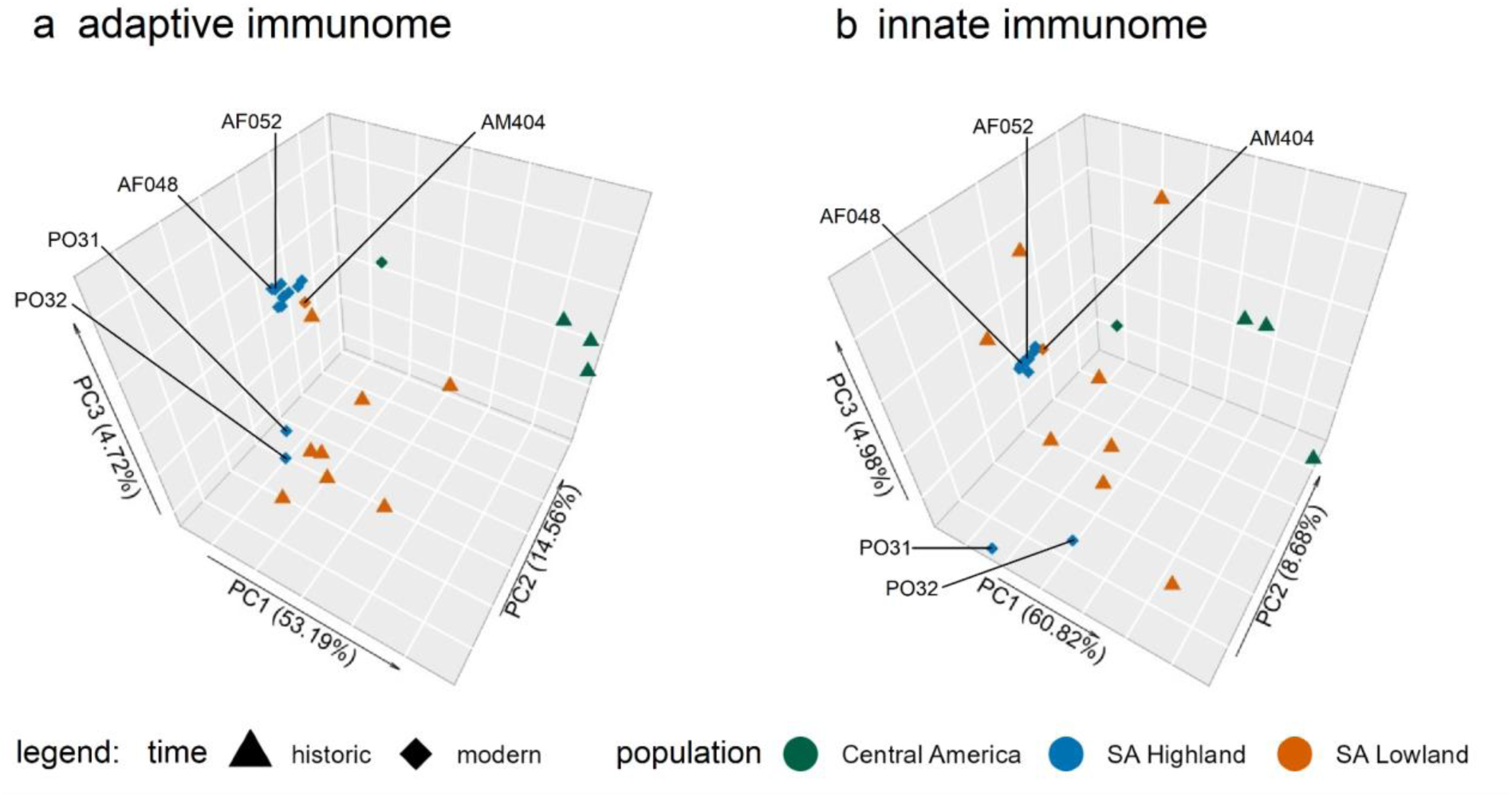
Principal component analysis of 26,408 unlinked adaptive immunome-wide SNPs (a), and principal component analysis of 14,155 unlinked innate immunome-wide SNPs (b) of 25 jaguar individuals including the first three principal components.

**SUPPLEMENTARY FIGURE S2:**
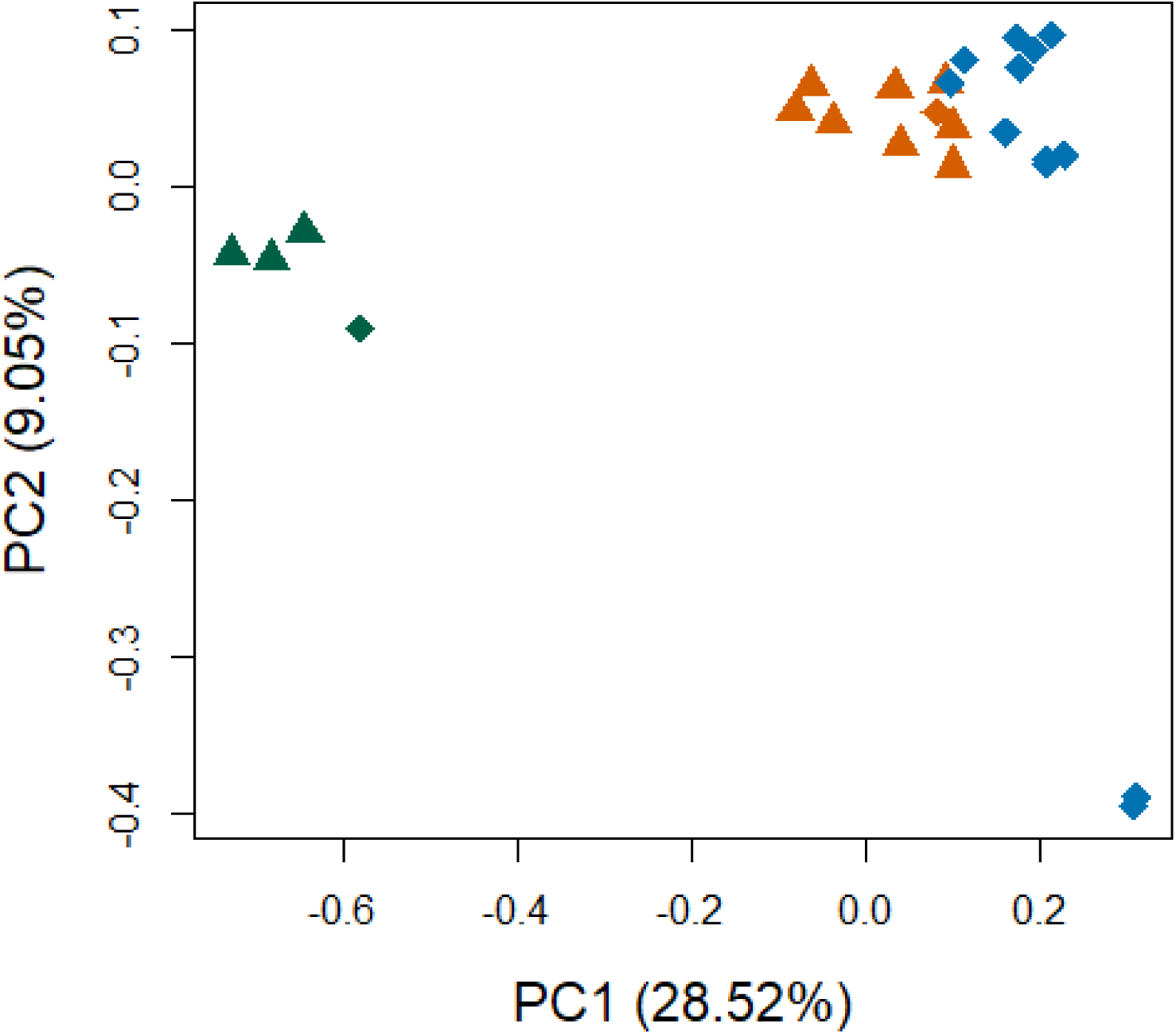
PCA of 689,785 unlinked genome-wide SNPs of 25 jaguar individuals showing the first two principal components, triangle represent historic samples, diamonds represent modern samples, green indicates samples from the Central American highland population, orange represents samples from the South American lowland population, blue represents samples from the South American highland population.

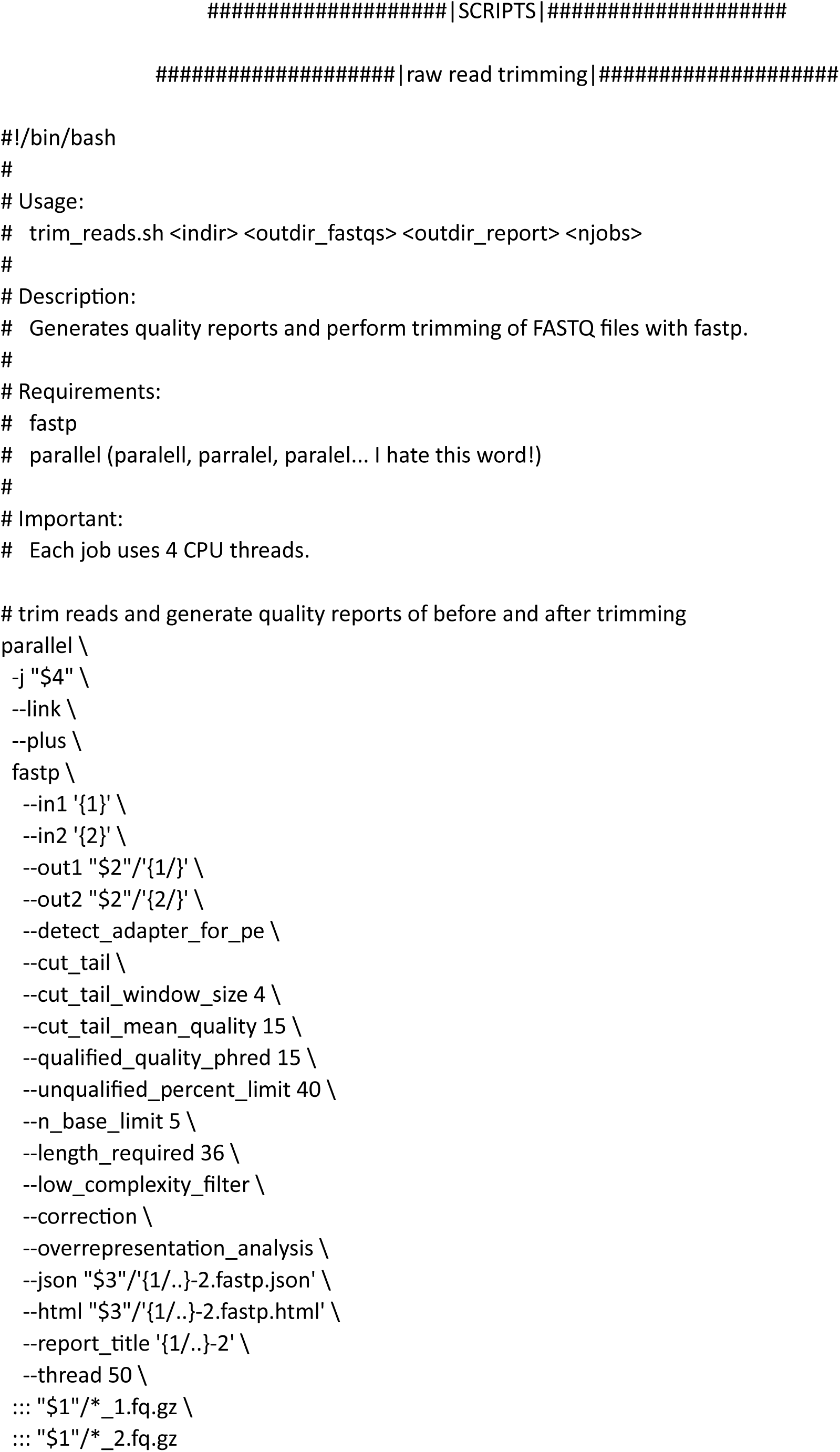
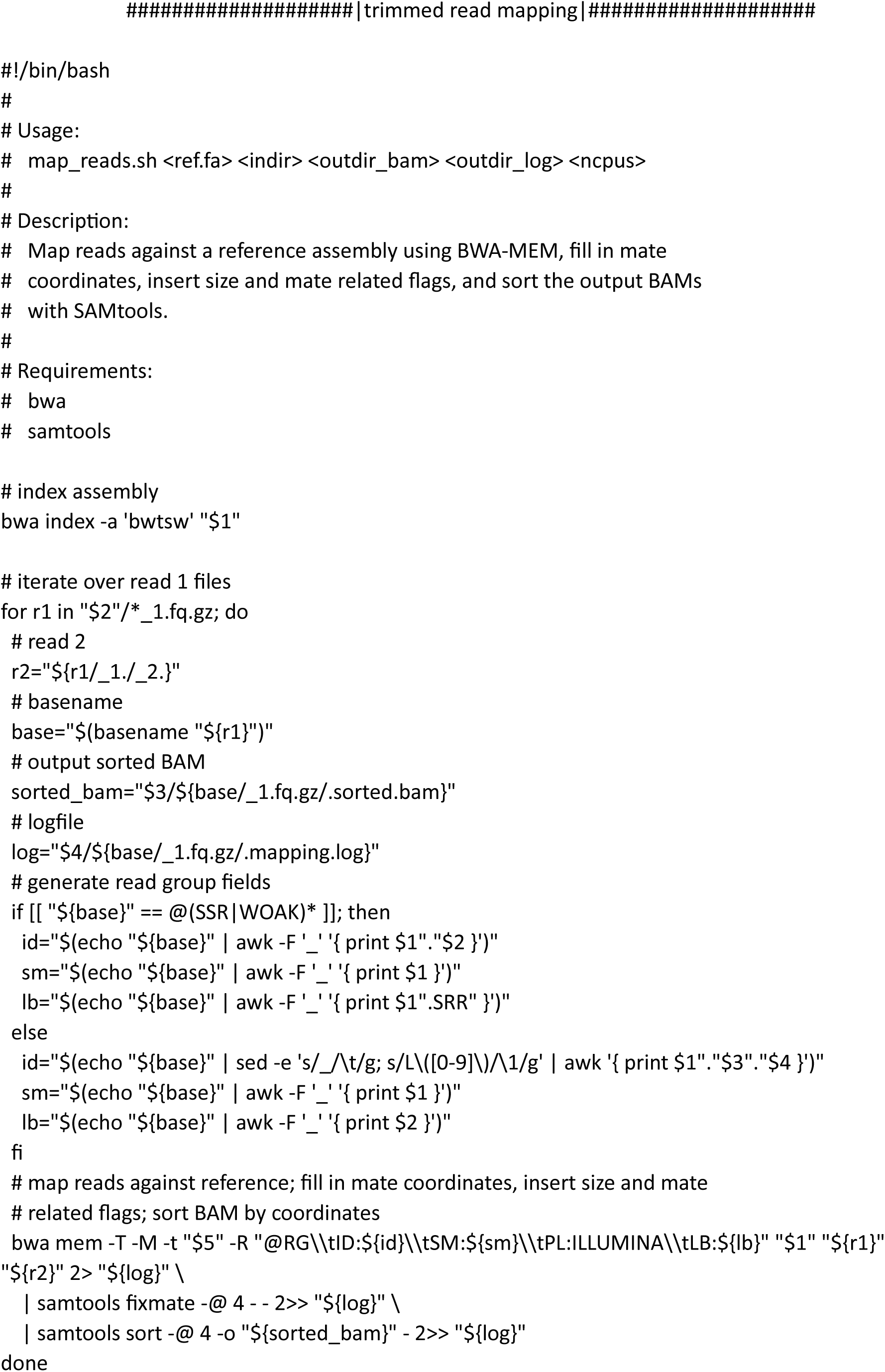
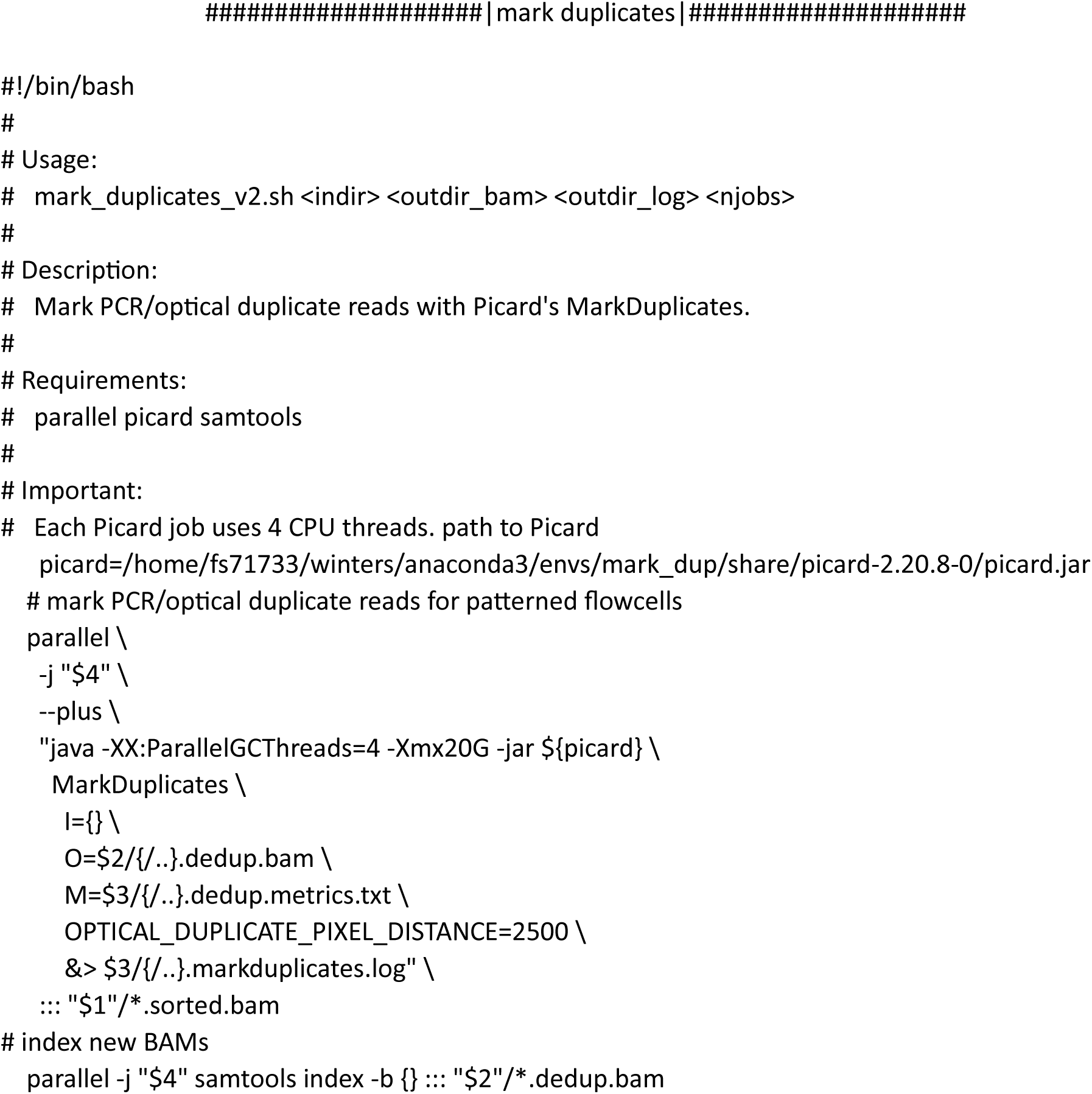
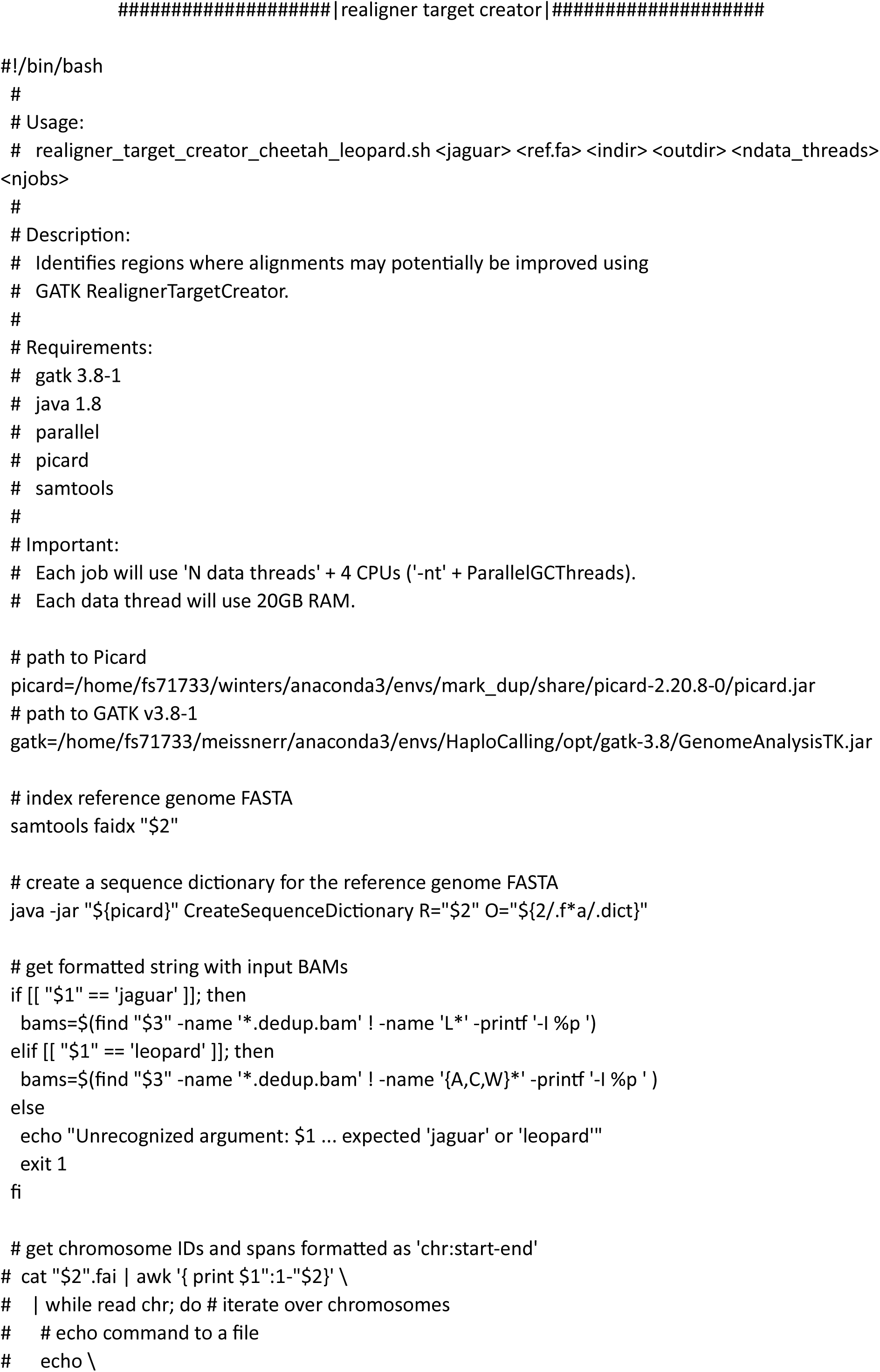
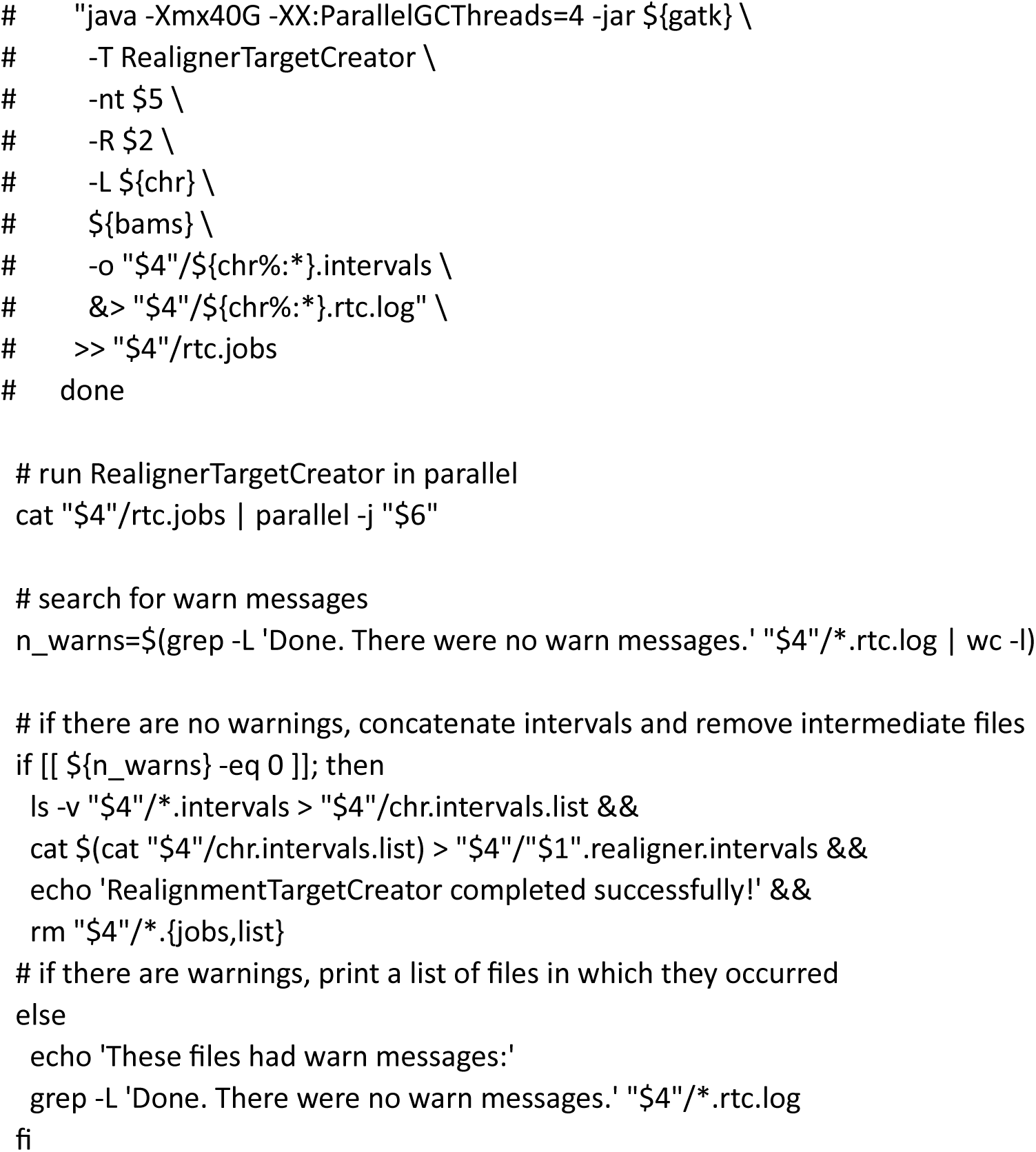
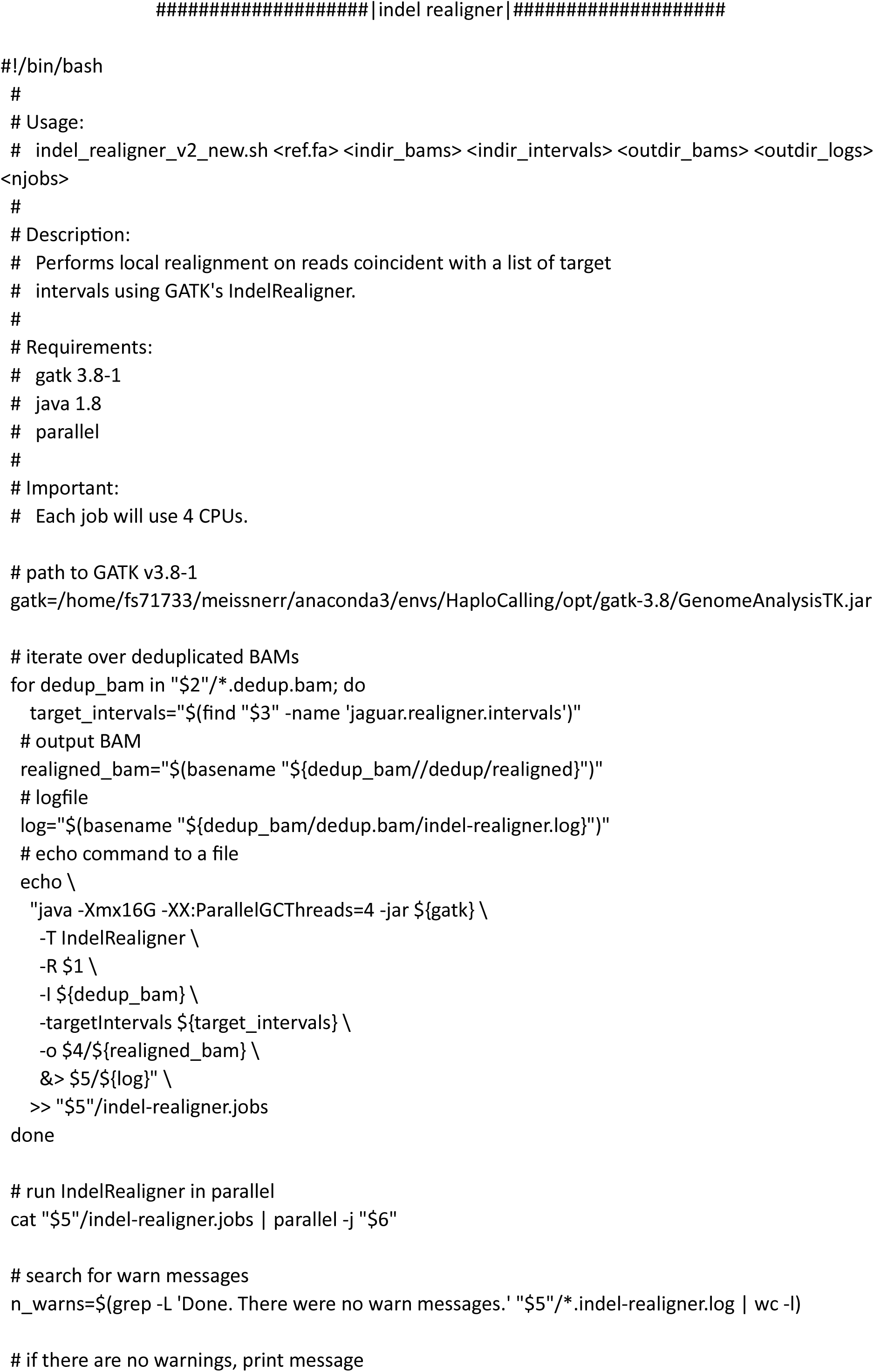
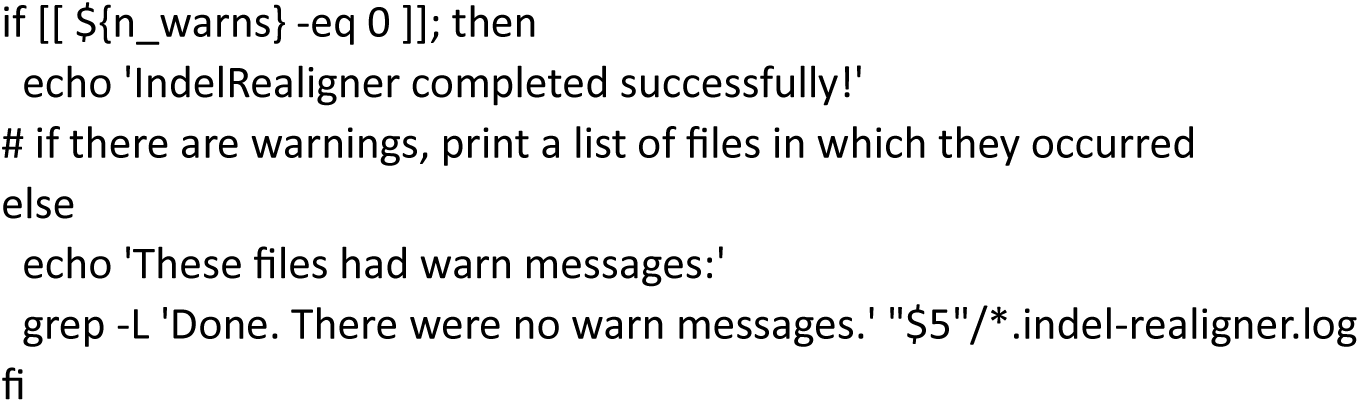
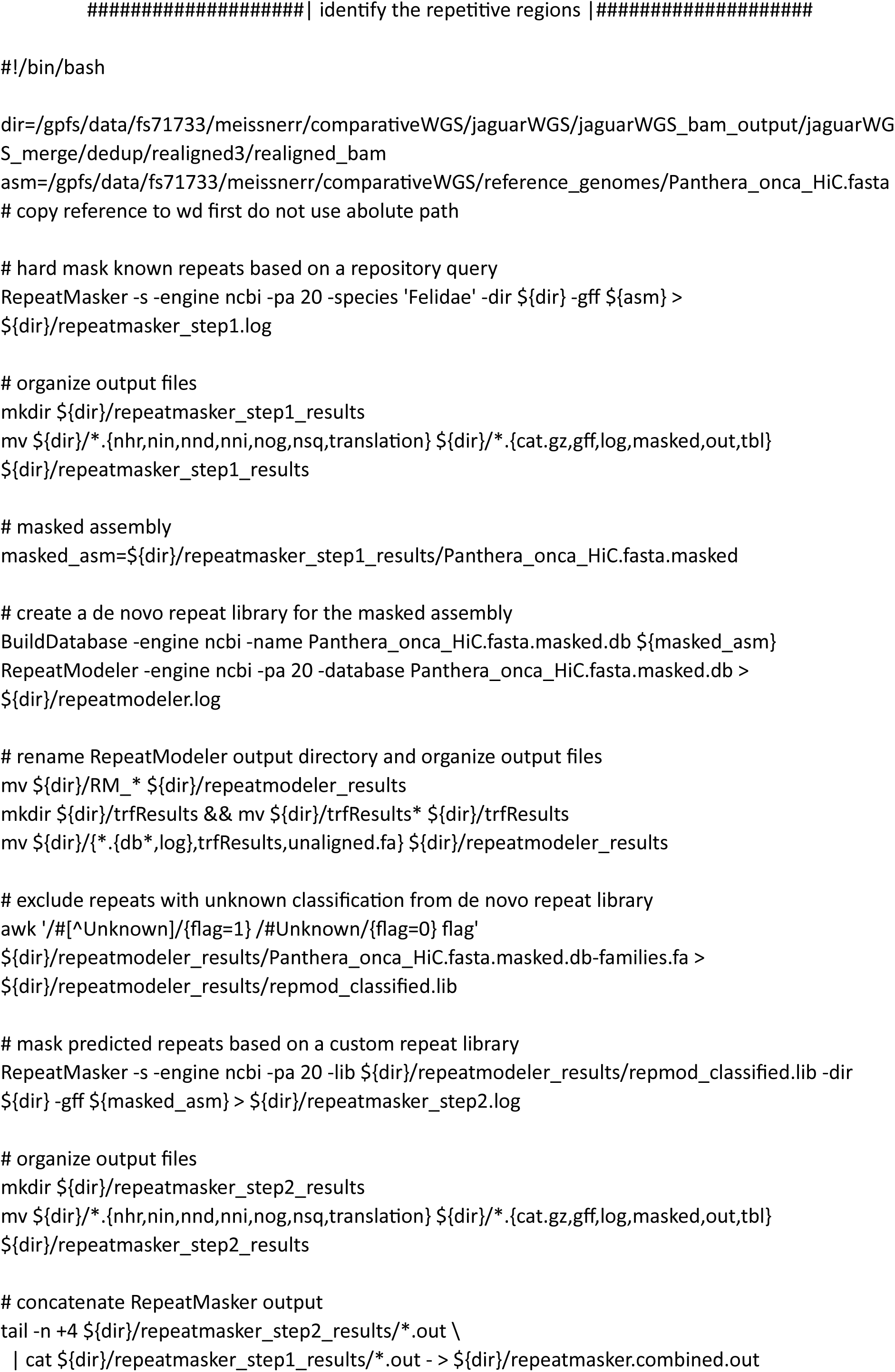
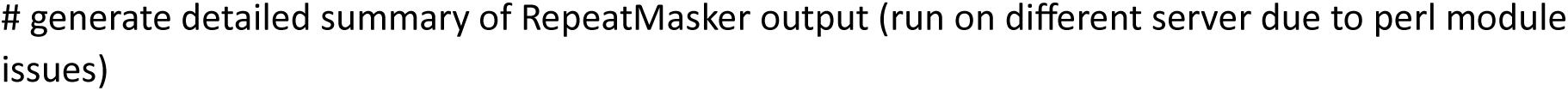
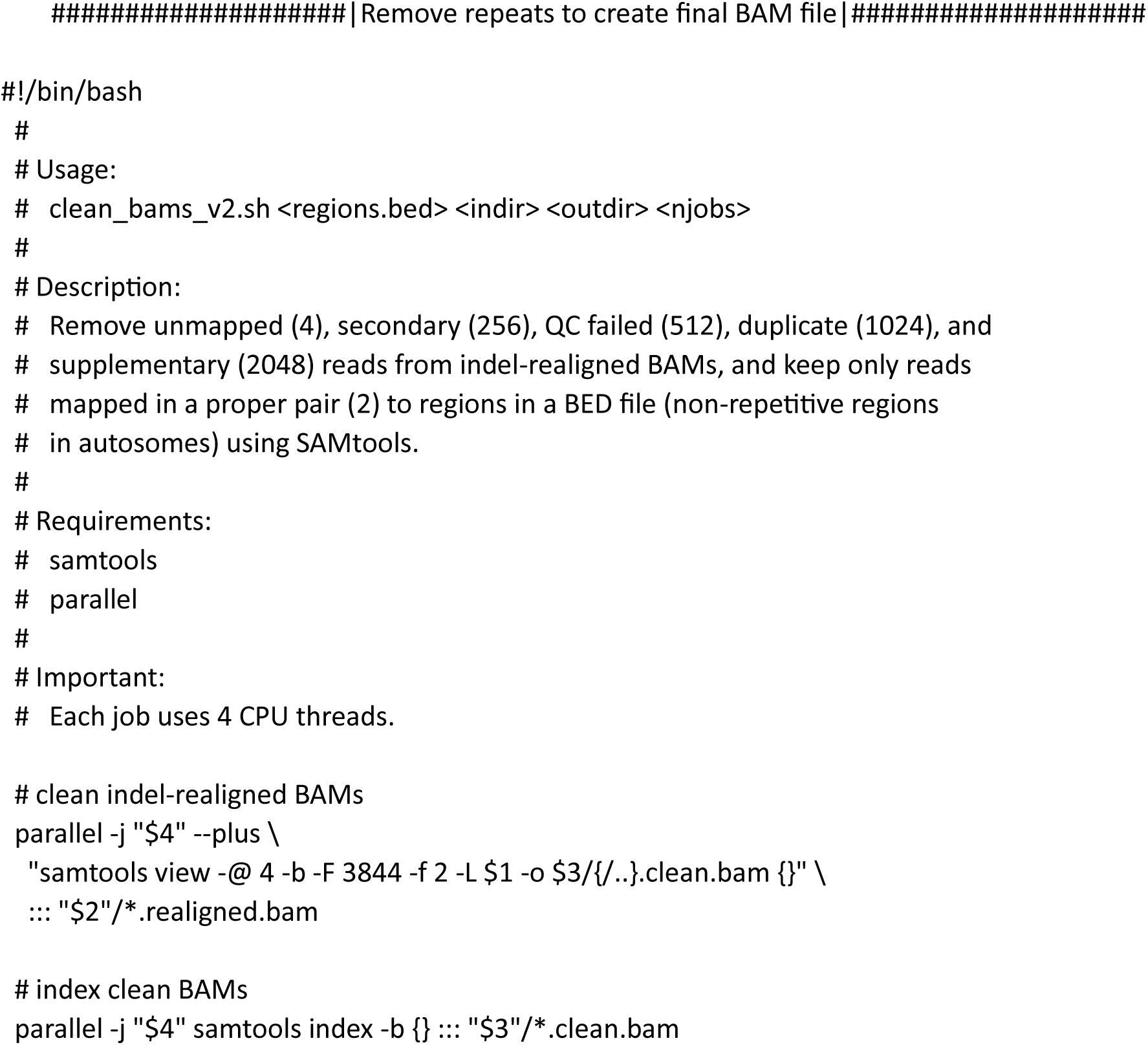
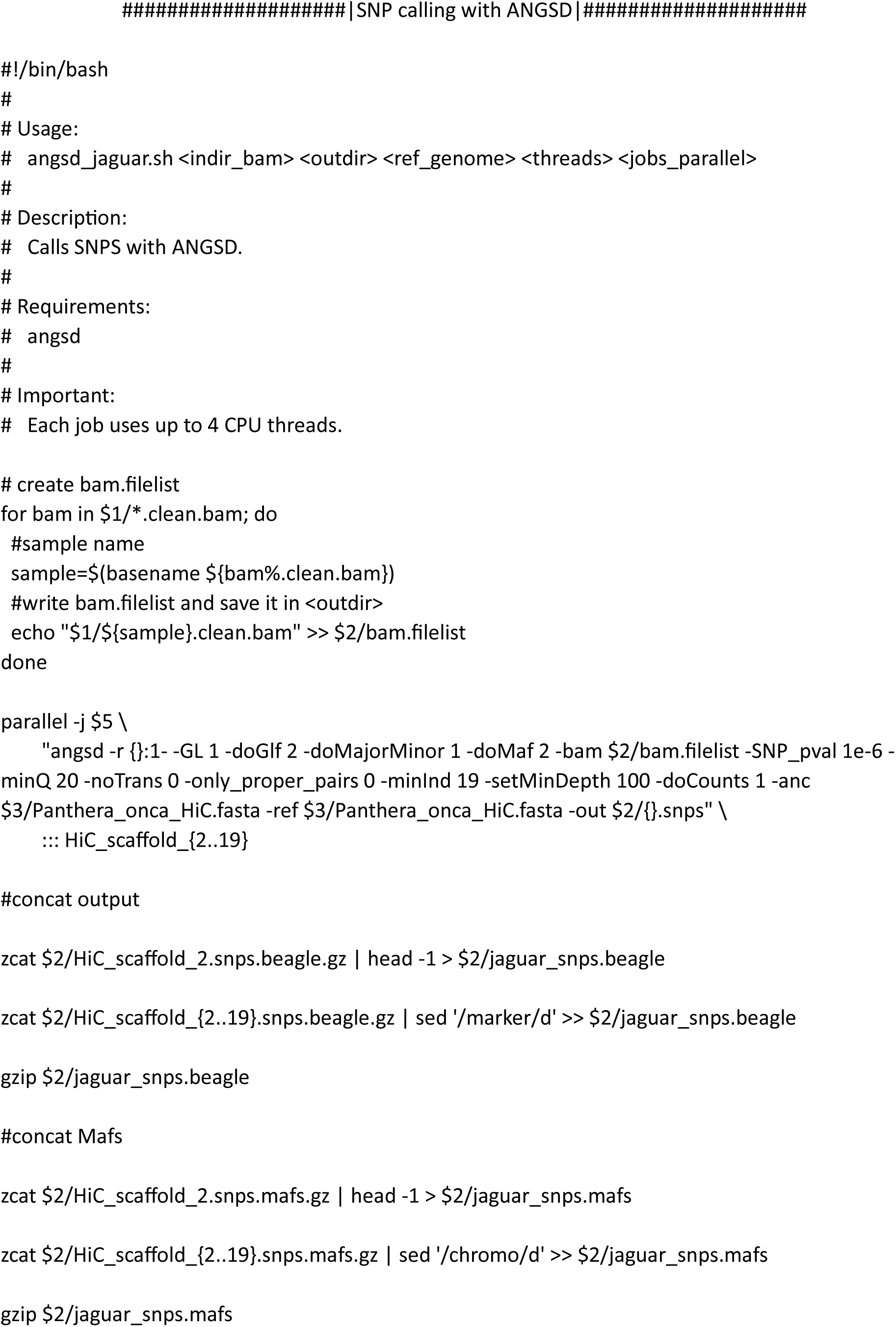
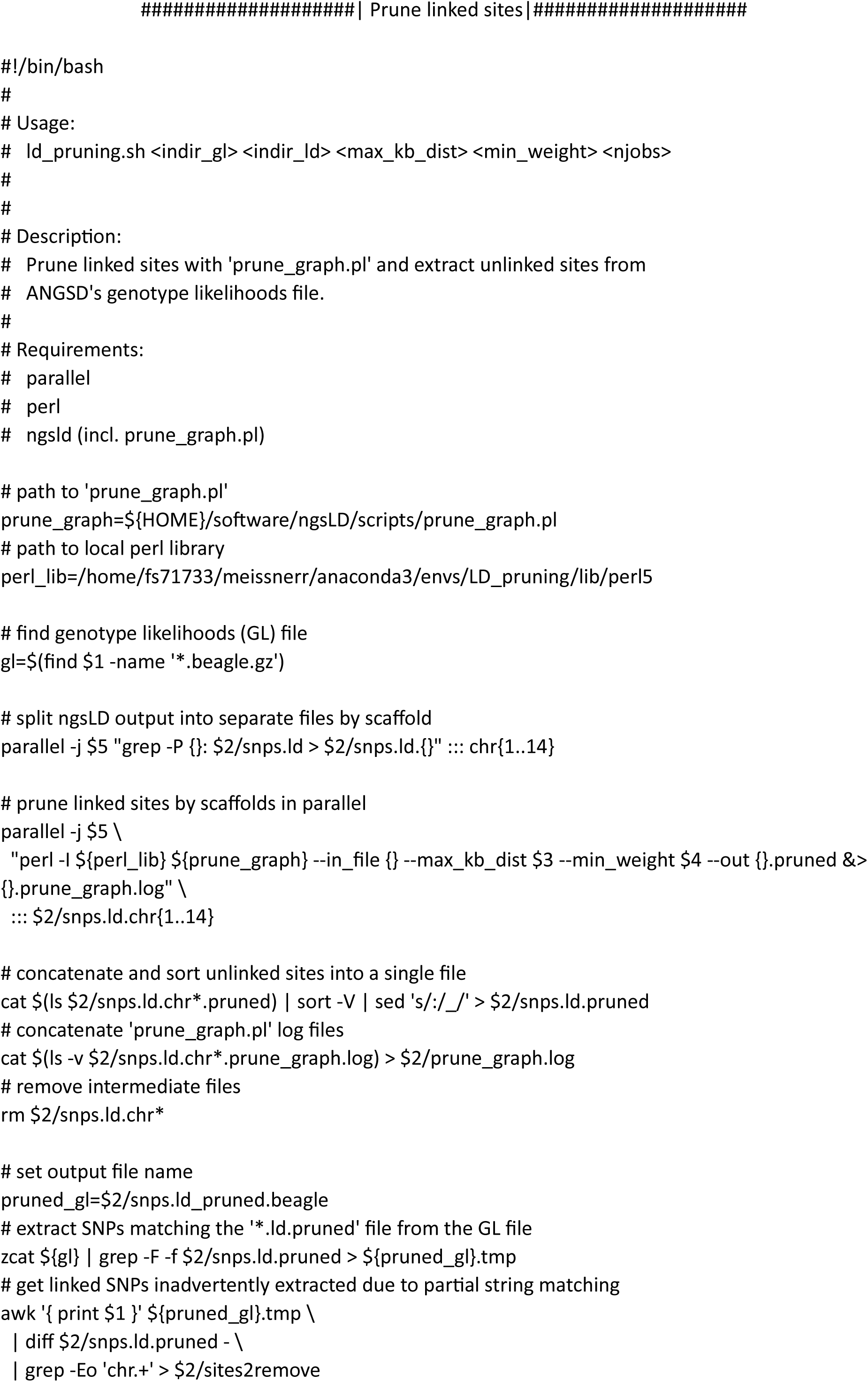
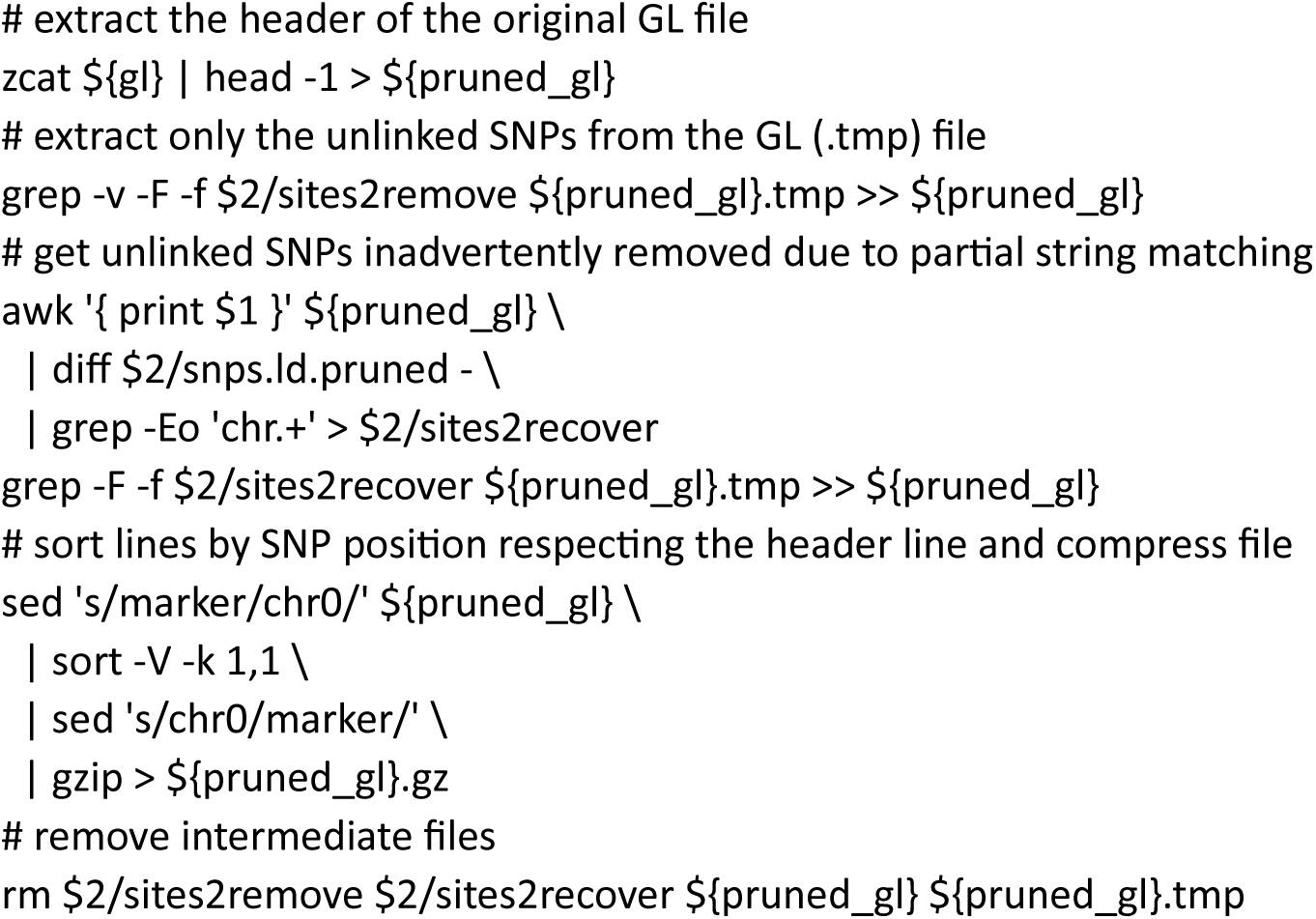
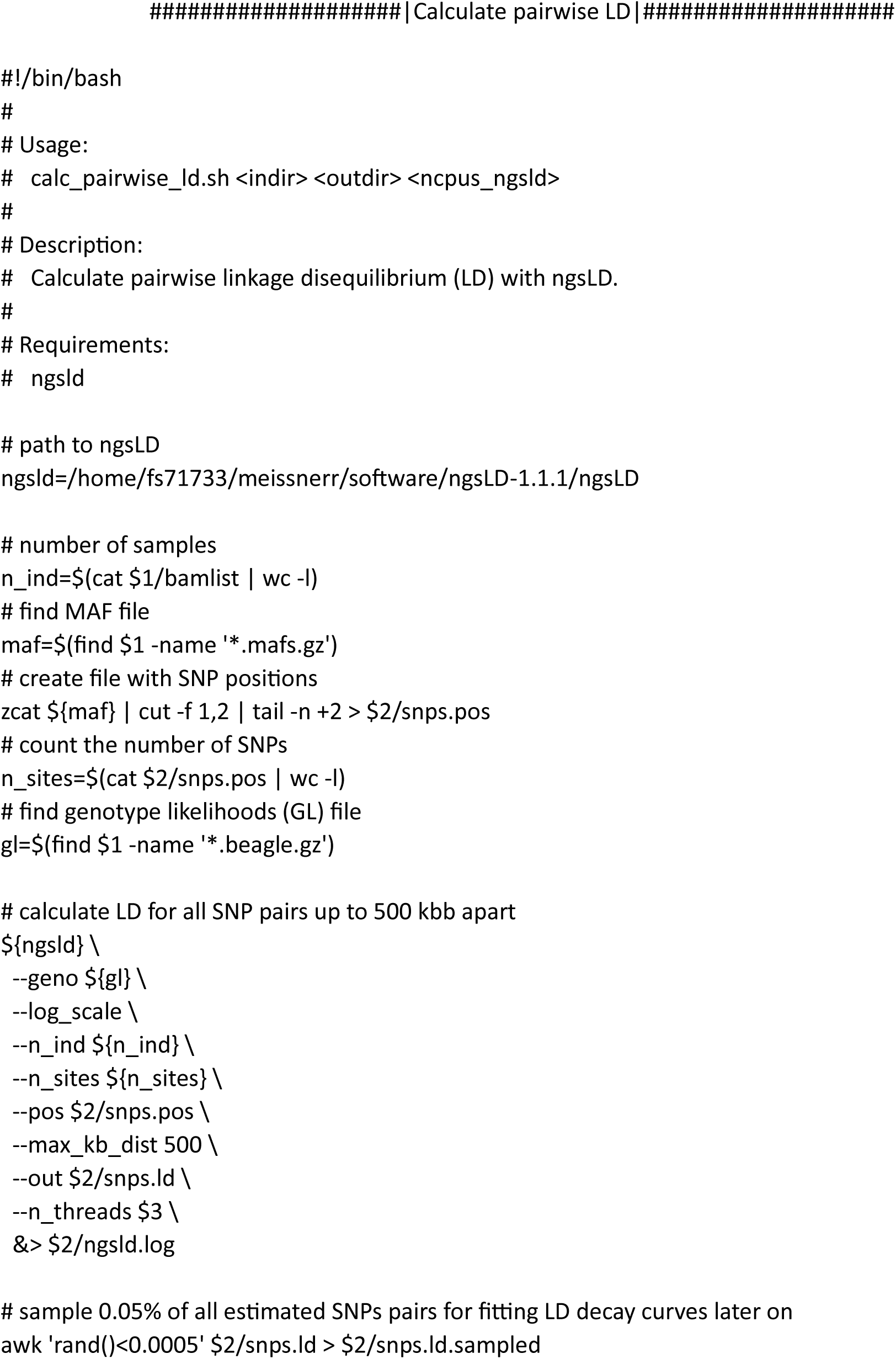
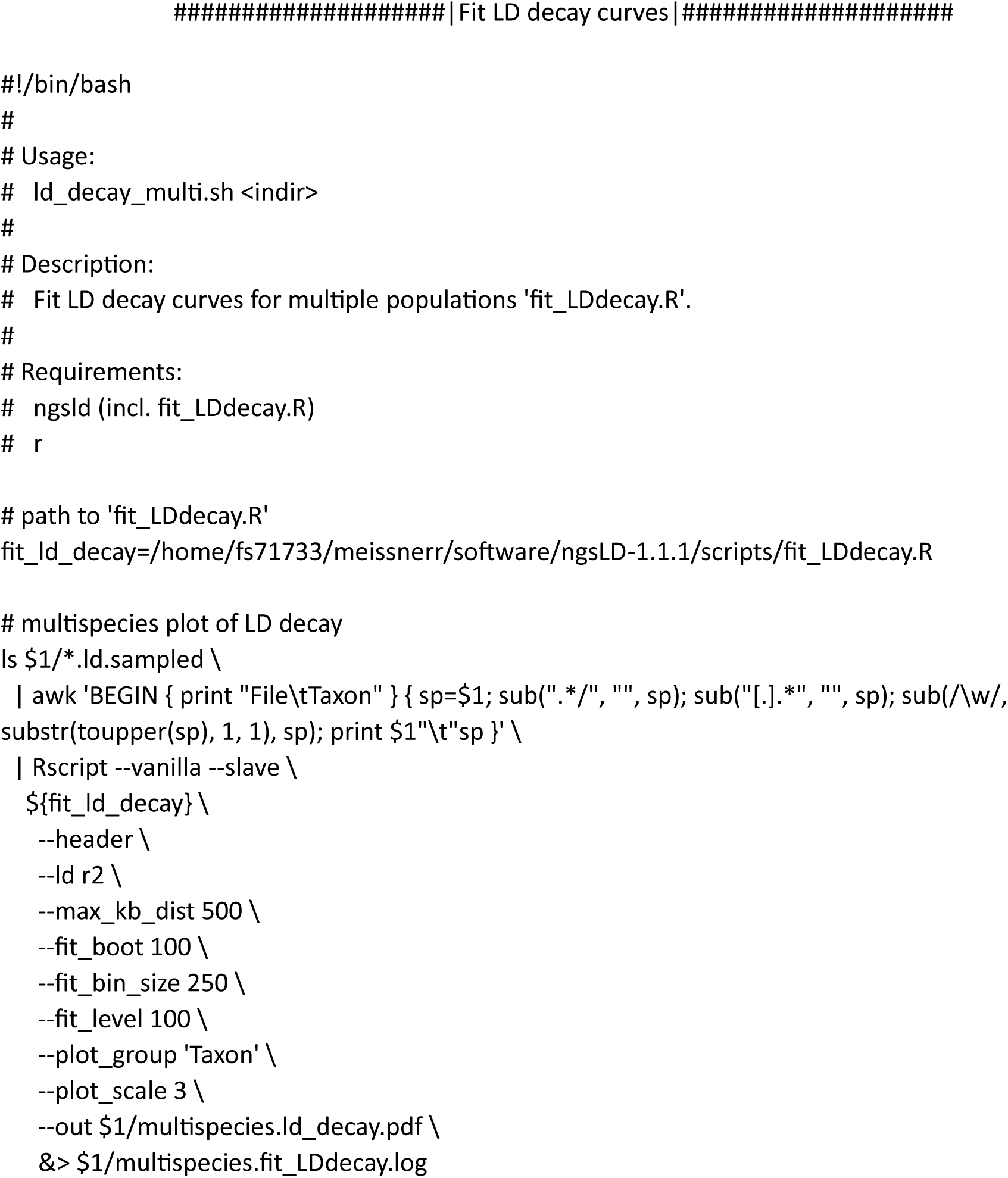
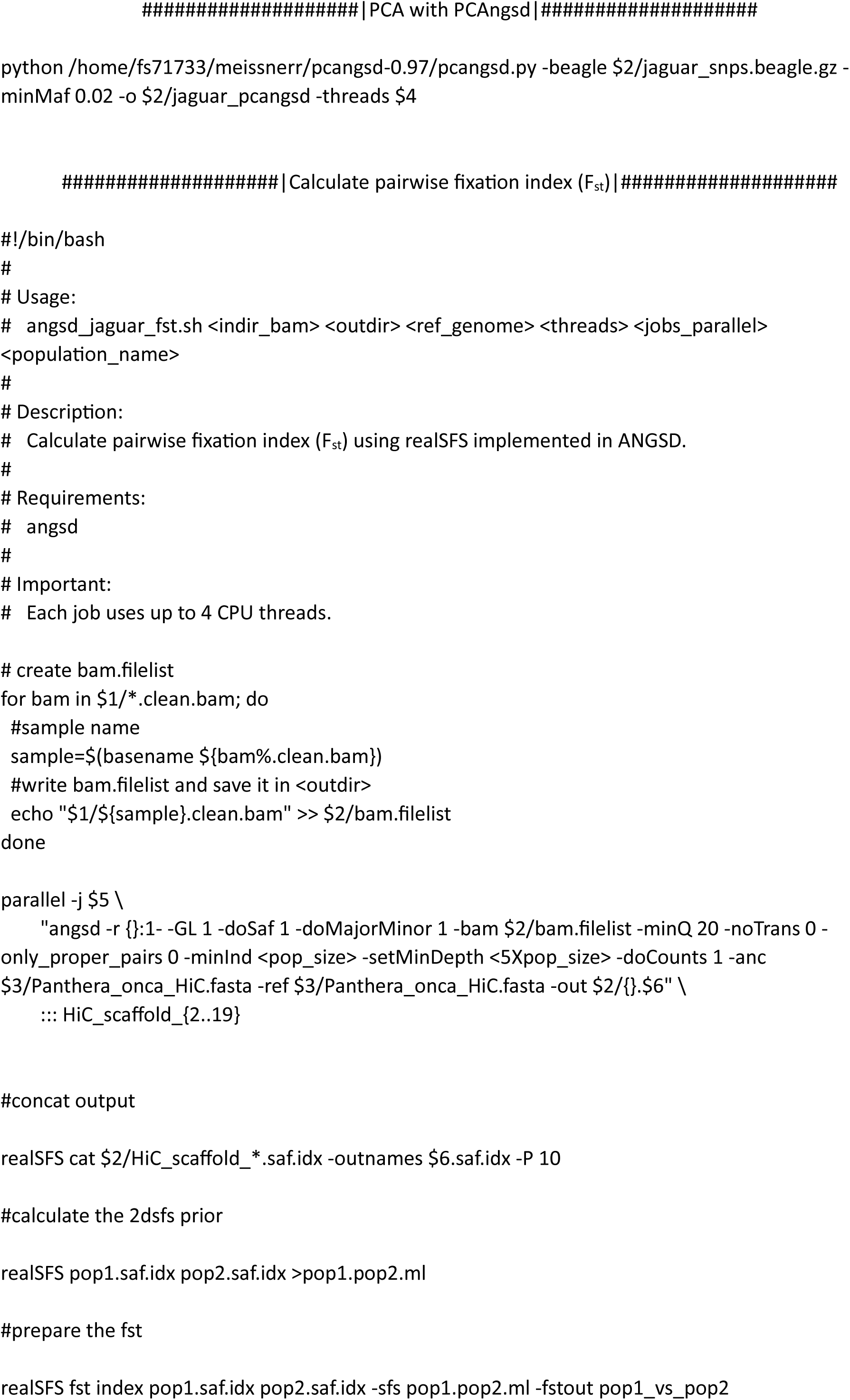
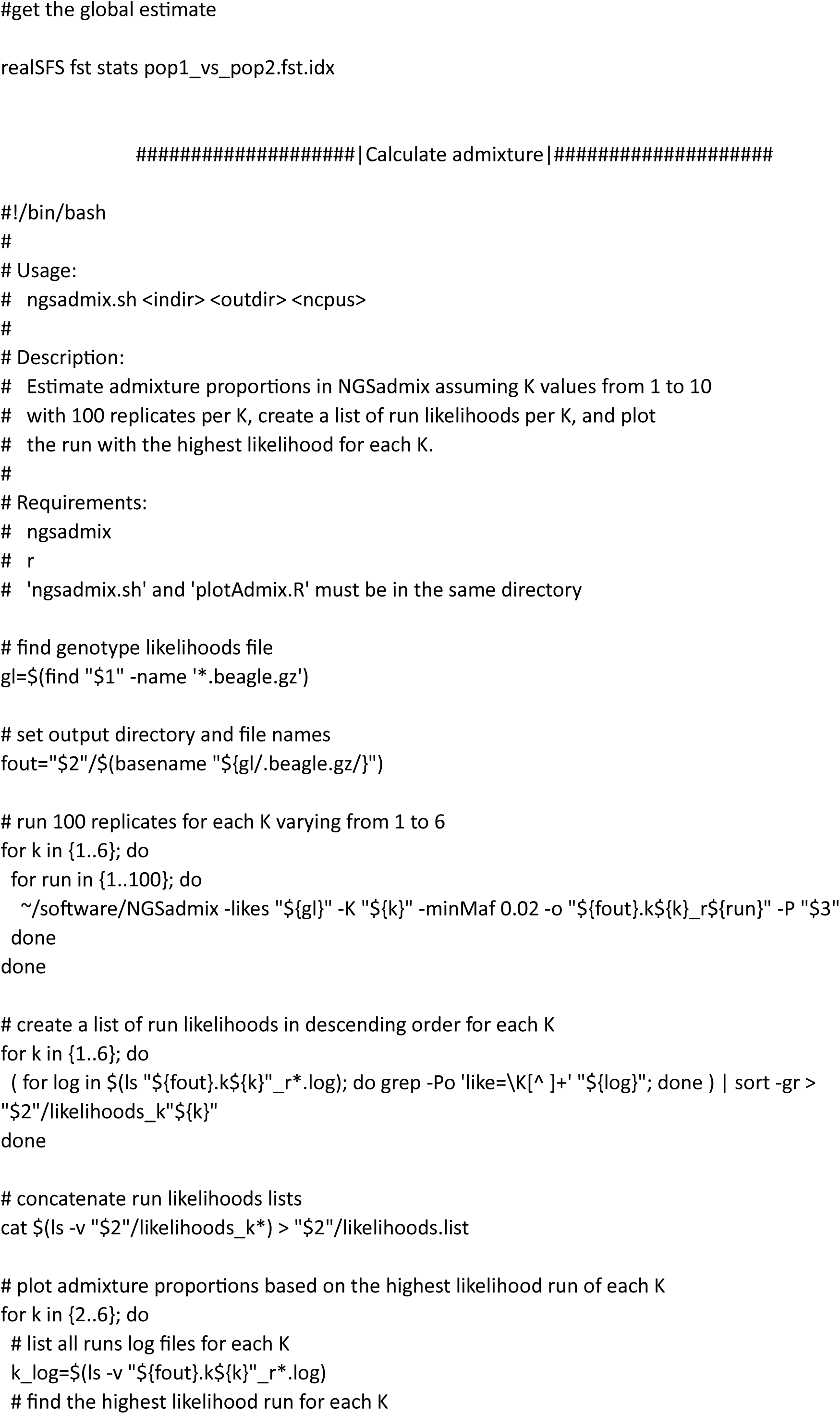
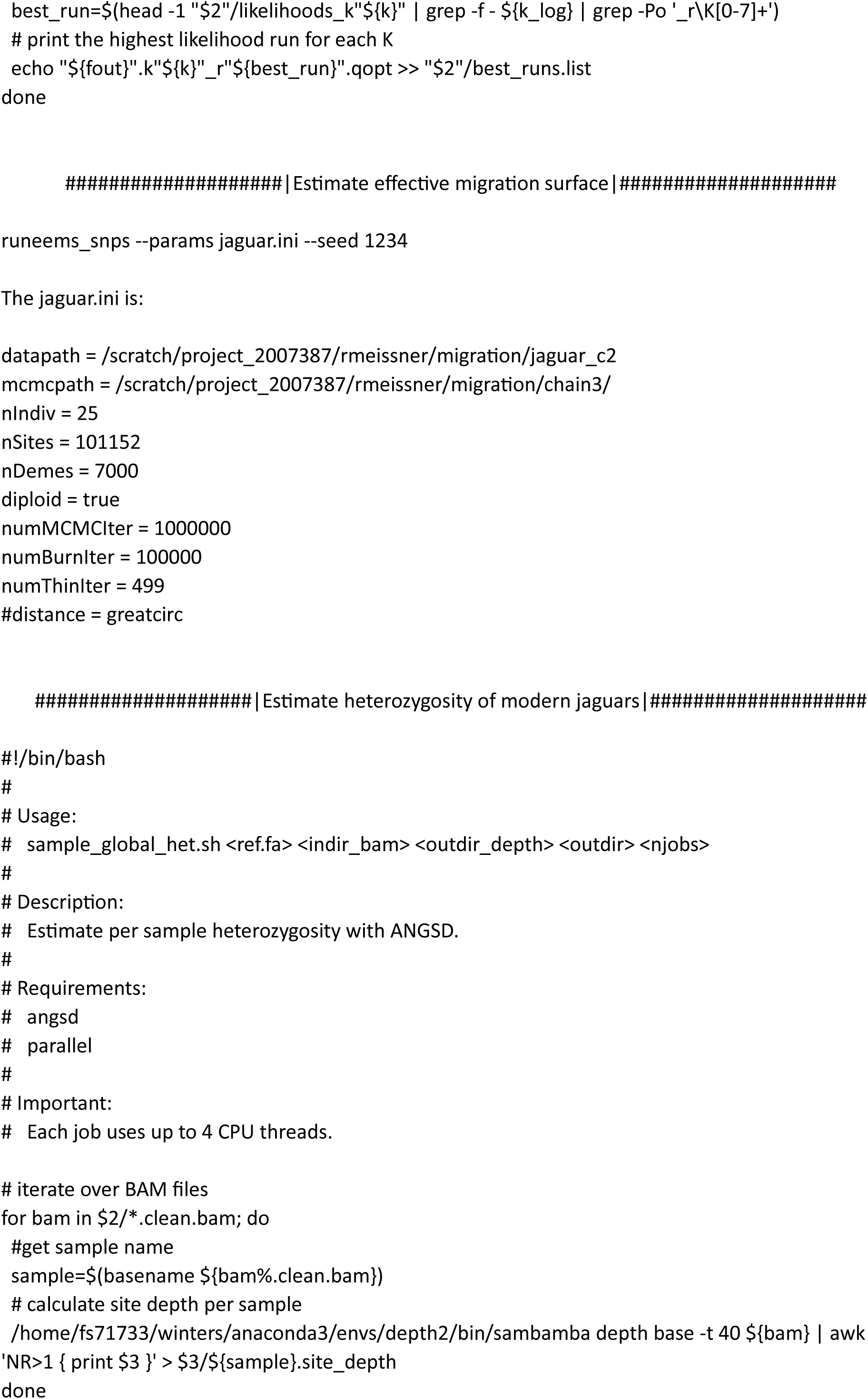
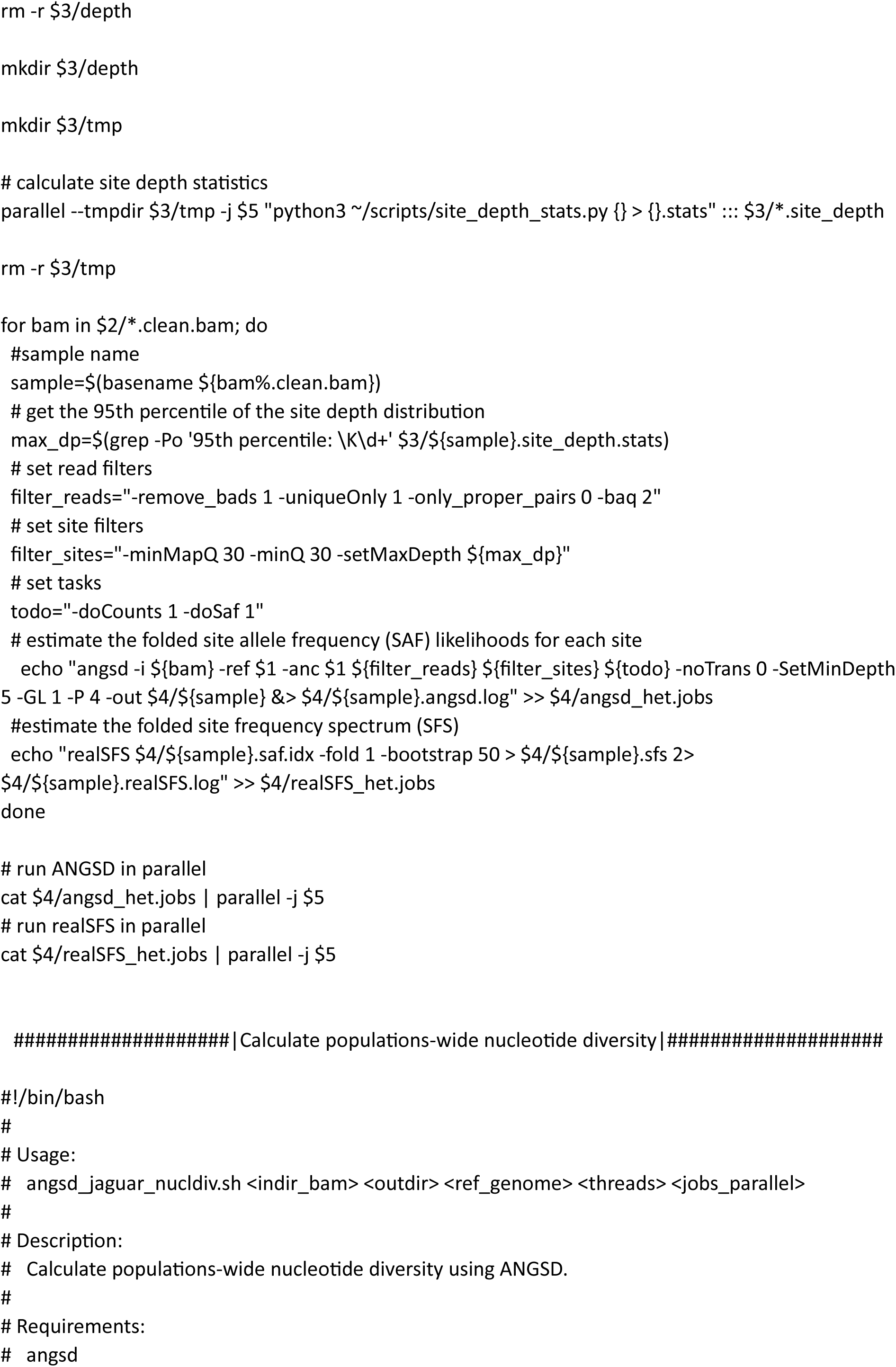
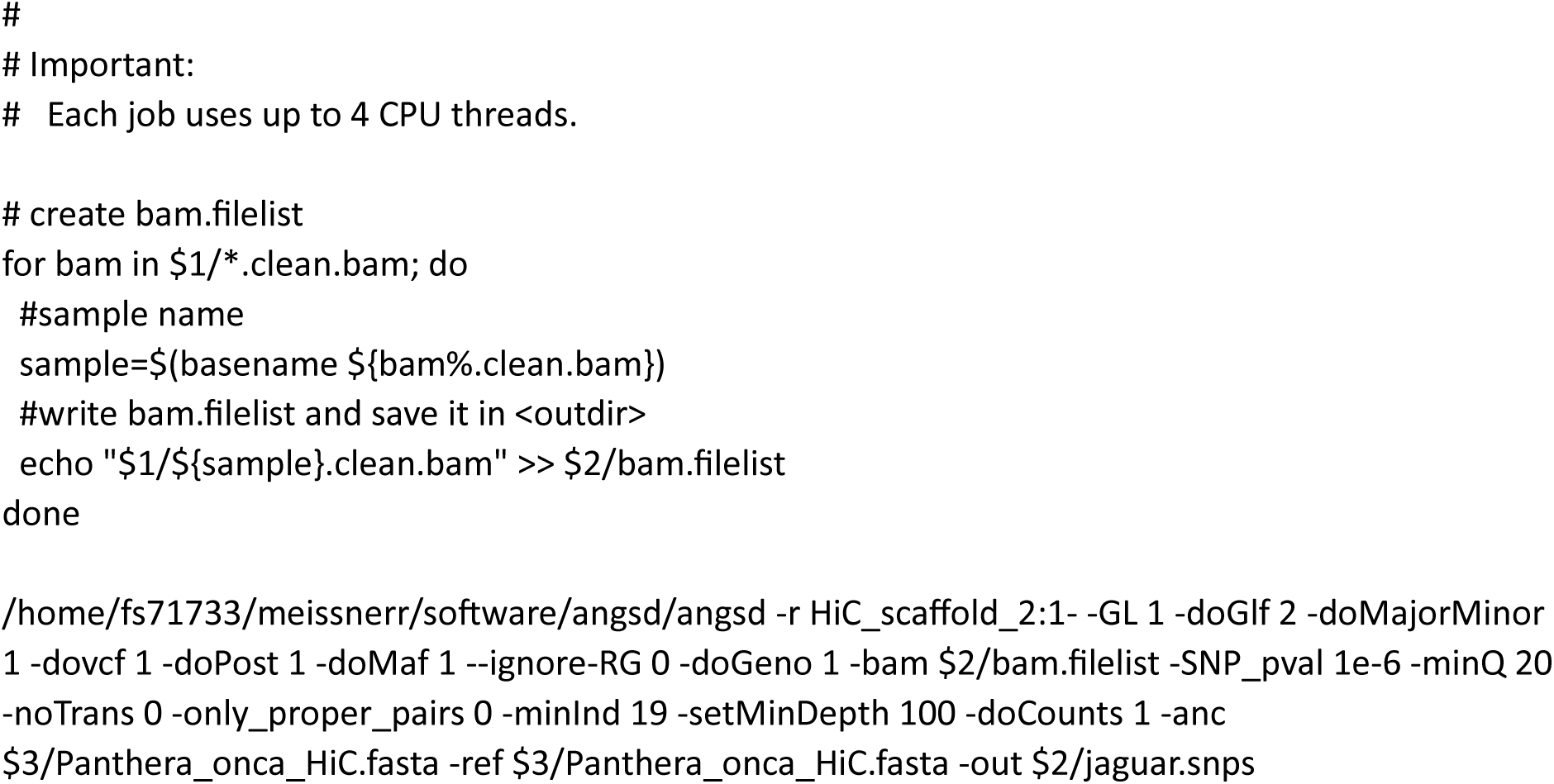
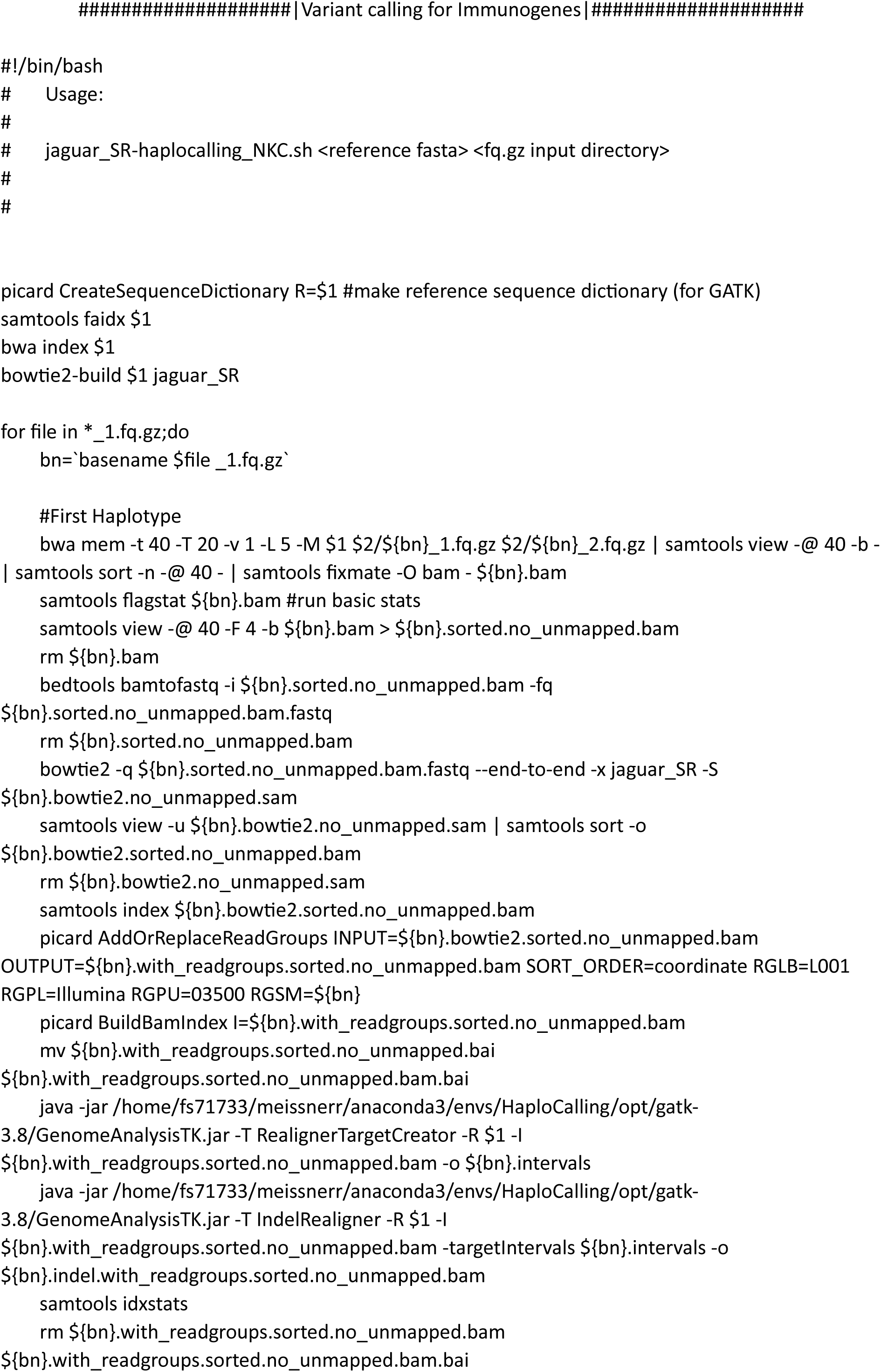
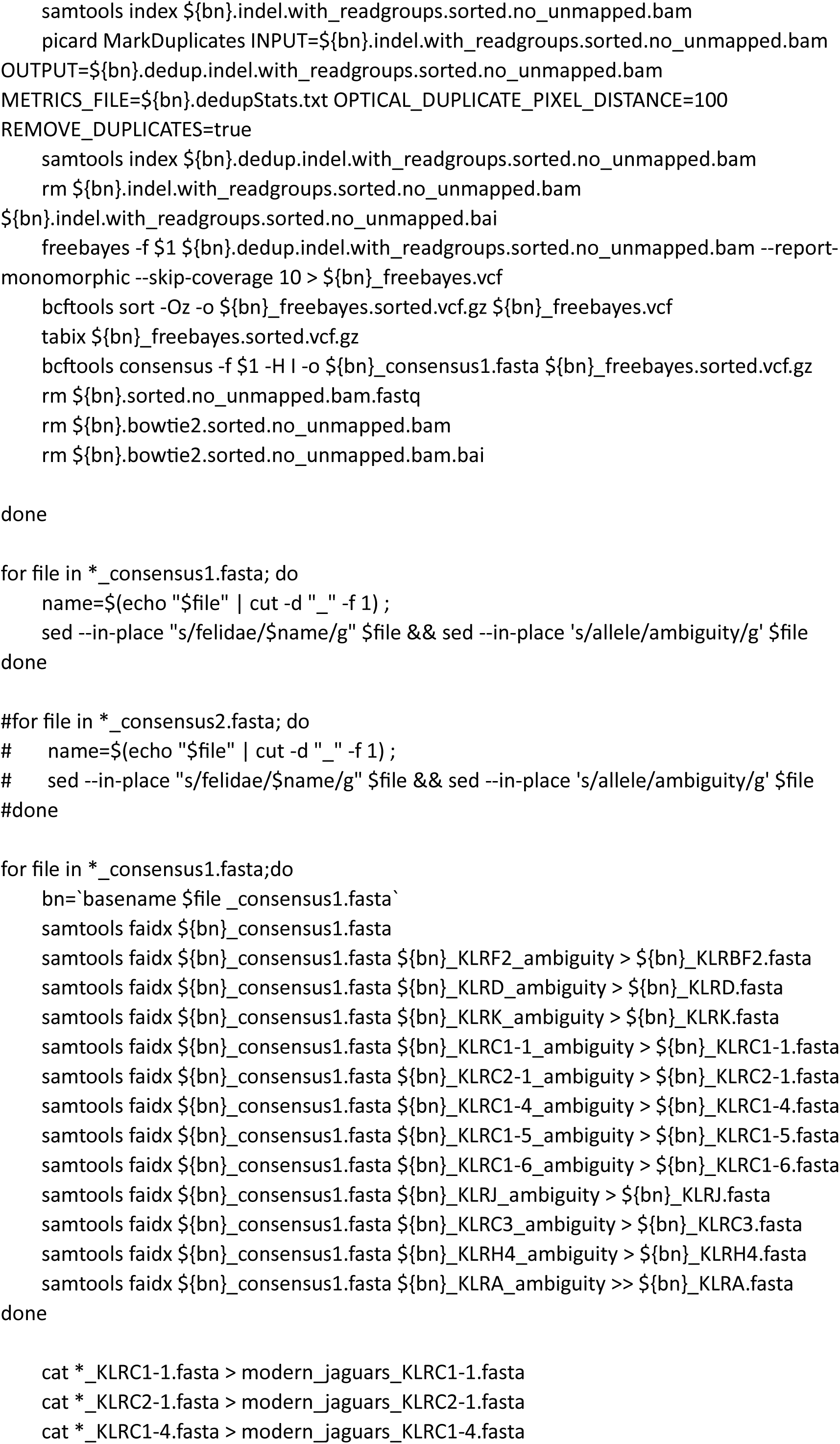
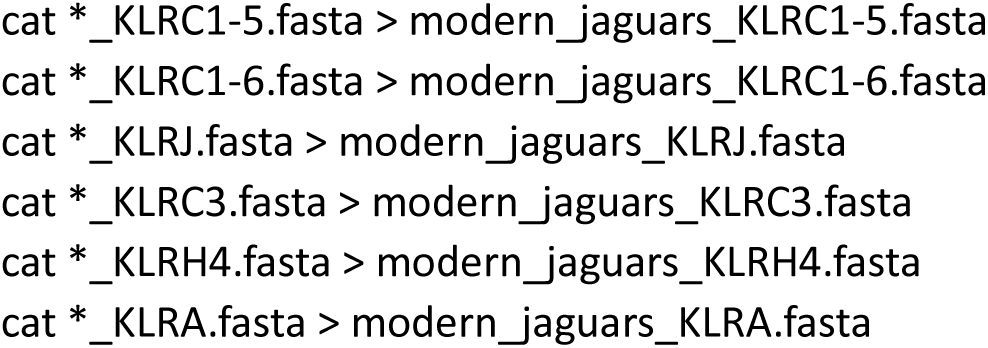

